# Targeting hypoxia-responsive miR-155-5p and miR-210-3p restores alveolar regeneration and reverses pulmonary fibrosis

**DOI:** 10.64898/2026.09.16.750970

**Authors:** Giulia Zandomenego, Alberto Maria Davide Ingo, Martina Torresi, Raffaella Klima, Maria Concetta Volpe, Francesco Salton, Paola Confalonieri, Karim Bahmed, Beata Kosmider, Ahmed A. Raslan, Danilo Licastro, Giovanni Ligresti, Marco Confalonieri, Luca Braga

**Affiliations:** Functional Cell Biology Group, International Centre for Genetic Engineering and Biotechnology (ICGEB), Trieste, Italy; Center for Inflammation and Lung Research, Lewis Katz School of Medicine,Temple University, Philadelphia, 19140, PA, United States; Department of Microbiology, Immunology and Inflammation, Lewis Katz School of Medicine, Temple University, Philadelphia, Pennsylvania, USA; Pulmonology Unit, Department of Medical Surgical and Health Sciences, Hospital of Cattinara, University of Trieste, 34149, Trieste, Italy; Boston University Pulmonary Centre, Chobanian and Avedisian School of Medicine, Boston, MA, USA; AREA Science Park, Trieste, Italy

## Abstract

Idiopathic pulmonary fibrosis (IPF) is an age-associated degenerative disease largely driven by failure of alveolar epithelial regeneration, yet current therapies slow fibrosis progression without restoring epithelial repair. Building on our previous unbiased microRNA screen in primary murine alveolar type II (ATII) cells, we identified miR-155-5p and miR-210-3p as previously unrecognized mediators of alveolar epithelial regenerative failure. Both microRNAs were markedly upregulated in ATII cells from patients with IPF and enriched within KRT17+/KRT5− aberrant transitional epithelial cells. Their expression also increased spontaneously with ageing in ATII cells from uninjured mice, linking these microRNAs to the age-dependent loss of ATII to ATI transdifferentiation capacity. Furthermore, we found that hypoxia-induced HIF signalling drives the expression of these microRNAs in ATII cells, locking them in a dysfunctional transitional state characterized by a profibrotic secretome that promotes paracrine myofibroblasts activation. Antisense oligonucleotide (ASO) inhibition of selected miRNAs restored ATII-to-ATI differentiation, eliminated aberrant transitional states, normalized epithelial-mesenchymal communication, promoted de novo alveolar regeneration and reversed fibrosis, including in both young and aged mice. Together, these findings identify the hypoxia-responsive miRNAs miR-155-5p and miR-210-3p as therapeutically actionable regulators of alveolar regenerative failure and establish their inhibition as a strategy to restore endogenous lung repair while disrupting pathological epithelial-mesenchymal crosstalk in pulmonary fibrosis.

## Introduction

Unlike most internal organs, the lung is continuously exposed to environmental insults, including pathogens, pollutants, mechanical stress and fluctuations in oxygen tension. Preserving the integrity of the gas-exchanging surface therefore depends on alveolar type II (ATII) cells, which secrete pulmonary surfactant and serve as facultative stem cells throughout life by self-renewing and differentiating following injury into alveolar type I (ATI) cells, the highly specialized epithelial cells that form the alveolar gas-exchange surface ^1–3^. However, ageing progressively compromises this regenerative competence through telomere attrition, mitochondrial dysfunction, epigenetic remodelling and cellular senescence. As regenerative capacity declines, physiological alveolar repair is progressively replaced by maladaptive fibrotic remodelling ^4^.

Idiopathic pulmonary fibrosis (IPF) is an age-associated degenerative lung disease characterized by progressive destruction of the alveolar architecture and irreversible respiratory failure. IPF is increasingly recognized as a disease of failed epithelial regeneration rather than simply uncontrolled fibroblasts activation ^5^.

Repetitive epithelial injury drives ATII cells into aberrant transitional states unable to complete differentiation into mature ATI cells ^6^. Single-cell transcriptomic analysis has identified these dysfunctional epithelial populations as KRT8⁺ transitional cells in mice ^7^ and KRT17⁺/KRT5⁻ basaloid cells in humans ^6^. Rather than supporting tissue repair, these cells acquire a maladaptive phenotype characterized by persistent stress responses, activation of developmental signalling pathways and secretion of profibrotic mediators that perpetuate fibroblast activation and extracellular matrix deposition ^8^. Progressive remodelling of the alveolar niche further impairs oxygen diffusion establishing chronic tissue hypoxia, a central amplifier of fibrotic remodelling ^9^. Although transient HIF activation contributes to physiological adaptation following injury ^10,11^, persistent HIF signalling can drive metabolic reprogramming and impair epithelial differentiation and regenerative capacity, thereby promoting maladaptive repair and fibrosis ^12^. Together, epithelial senescence, maladaptive transitional cell states and chronic hypoxic signalling have emerged as convergent mechanisms underlying disease progression.

Despite this growing mechanistic understanding, currently approved therapies remain limited. Pirfenidone, nintedanib and, more recently, nerandomilast slow disease progression but have not been shown to restore endogenous alveolar regeneration ^13–15^.

Moreover, the repeated failure of late-stage clinical trials targeting individual signalling pathways underscores the profound biological heterogeneity of IPF ^16^ ^17^ and suggests that successful therapies will need to restore coordinated regenerative programs rather than inhibit isolated downstream profibrotic mechanisms.

MicroRNAs (miRNAs) provide a potential means of achieving such coordinated regulation. By simultaneously modulating multiple transcripts and interconnected signalling pathways, miRNAs can orchestrate complex cellular states ^18^. Numerous miRNAs have been implicated in IPF, particularly through regulation of fibroblast activation, extracellular matrix deposition and epithelial-to-mesenchymal transition ^19–28^. However, despite the central role of alveolar epithelial dysfunction in IPF, only miR-375 ^29^, miR-200 ^30,31^ and, more recently by our group, miR-124-3p have been directly linked to ATII-to-ATI differentiation and alveolar repair ^32^. Therapeutic restoration of miR-124-3p through AAV6.2FF-mediated delivery showed efficacy in preclinical fibrosis models ^32^, although viral vector-based strategies remain constrained by manufacturing complexity, immunogenicity, limited re-administration feasibility and unresolved long-term safety concerns ^33^.

In contrast, antisense oligonucleotides (ASOs) have emerged as a clinically validated platform for therapeutic modulation of pathogenic RNA networks ^34^Krutzfeldt, 2005 #17}. Multiple ASO drugs targeting protein-coding transcripts have been approved for human disease ^35–38^, while anti-miRNA strategies have progressed from preclinical studies ^39–41^ to early-phase clinical trials ^42,43^, establishing the feasibility and translational potential of inhibitory RNA therapeutics, although efficacy in later-phase trials remains to be established ^28^. Their compatibility with repeat dosing, favourable pharmacological properties and established clinical development pathways make ASOs particularly attractive for chronic respiratory diseases ^34^.

A major unresolved question is therefore whether disease-driving epithelial miRNAs exist and, if so, whether they can coordinately regulate key pathological features of IPF, including hypoxia, epithelial senescence, ATII-to-ATI transitional cell-state arrest and defective regeneration, and whether therapeutic inhibition of these regulators can restore endogenous alveolar repair while simultaneously interrupting pathological epithelial–mesenchymal crosstalk.

Here, we identify miR-155-5p and miR-210-3p as hypoxia-responsive regulators of alveolar epithelial regenerative failure in IPF. Both miRNAs were increased in ATII cells from patients with IPF and enriched in KRT17⁺/KRT5⁻ aberrant transitional epithelial cells, providing direct evidence of their association with the dysfunctional human alveolar epithelium.

Across murine models in both young and aged mice, we further show that chronic HIF signalling induces miR-155-5p and miR-210-3p expression, coupling chronic hypoxia to epithelial senescence, regenerative arrest and pathological epithelial-mesenchymal communication. Mechanistically, hypoxic signalling drives miR-155-5p and miR-210-3p overexpression in ATII cells, which in turn converts otherwise healthy ATII cells into dysfunctional cells that fail to support alveolar repair while becoming profibrotic signalling hubs whose secretome is sufficient to activate neighbouring fibroblasts. Finally, therapeutic inhibition of either miRNA using ASOs reverses established fibrosis across multiple preclinical IPF models, including bleomycin-induced lung fibrosis in aged mice. Together, these findings identify a hypoxia-responsive epithelial miRNA network as a central regulator of regenerative failure and establish anti-miRNA therapy as a promising regenerative strategy for IPF.

## Results

### Phenotypic screening uncovers miR-155-5p and miR-210-3p as drivers of epithelial dysfunction in IPF

We hypothesized that microRNAs simultaneously impairing epithelial differentiation and inducing senescence could act as key regulators of epithelial regenerative failure in IPF ^8^ ^44^ and therefore represent attractive therapeutic targets. To identify such regulators, we performed an analysis of a previously generated in vitro screen using human miRNA mimics in primary ATII cells ^32^. We specifically interrogated miRNAs that produced a combined phenotype of impaired ATII-to-ATI differentiation, measured by reduced numbers of RAGE⁺ cells, and increased cellular senescence, assessed by nuclear accumulation of p21. By integrating these complementary readouts, we sought to identify miRNAs capable of simultaneously disrupting alveolar epithelial differentiation and promoting a senescent cell state, two key features of epithelial dysfunction in IPF.

Among the 2,042 human microRNAs screened, only three microRNAs: hsa-miR-155-5p, hsa-miR-210-3p, and hsa-miR-490-3p recapitulate this pathogenic phenotype in healthy ATII cells (Fig. 1a and Extended Data Fig. 1a-b). All these three microRNAs are evolutionary conserved between human and mouse and represent potential candidates amenable to therapeutic inhibition. We next assessed their expression in the bleomycin-induced mouse model of pulmonary fibrosis, 21 days after bleomycin administration, corresponding to a phase of established fibrosis ^5^. Only miR-155-5p and miR-210-3p were significantly upregulated at this timepoint, whereas miR-490-3p remained minimally expressed and was not induced following bleomycin injury (Extended Data Fig. 1c-f).

We therefore focused on miR-155-5p and miR-210-3p and assessed their clinical relevance in human IPF lungs (Fig. 1b). miRNA-FISH combined with SPC staining revealed a marked increase in miR-155-5p in IPF lungs, with an approximately 3.3-fold higher proportion of positive cells than in healthy lungs (Fig. 1c,d). Consistently, mature miR-155-5p levels were significantly increased in ATII cells isolated from patients with IPF (Fig. 1e). A similar pattern was observed for miR-210-3p, which showed increased abundance in IPF lungs and in isolated IPF ATII cells (Fig. 1f–h).

**Figure 1.**
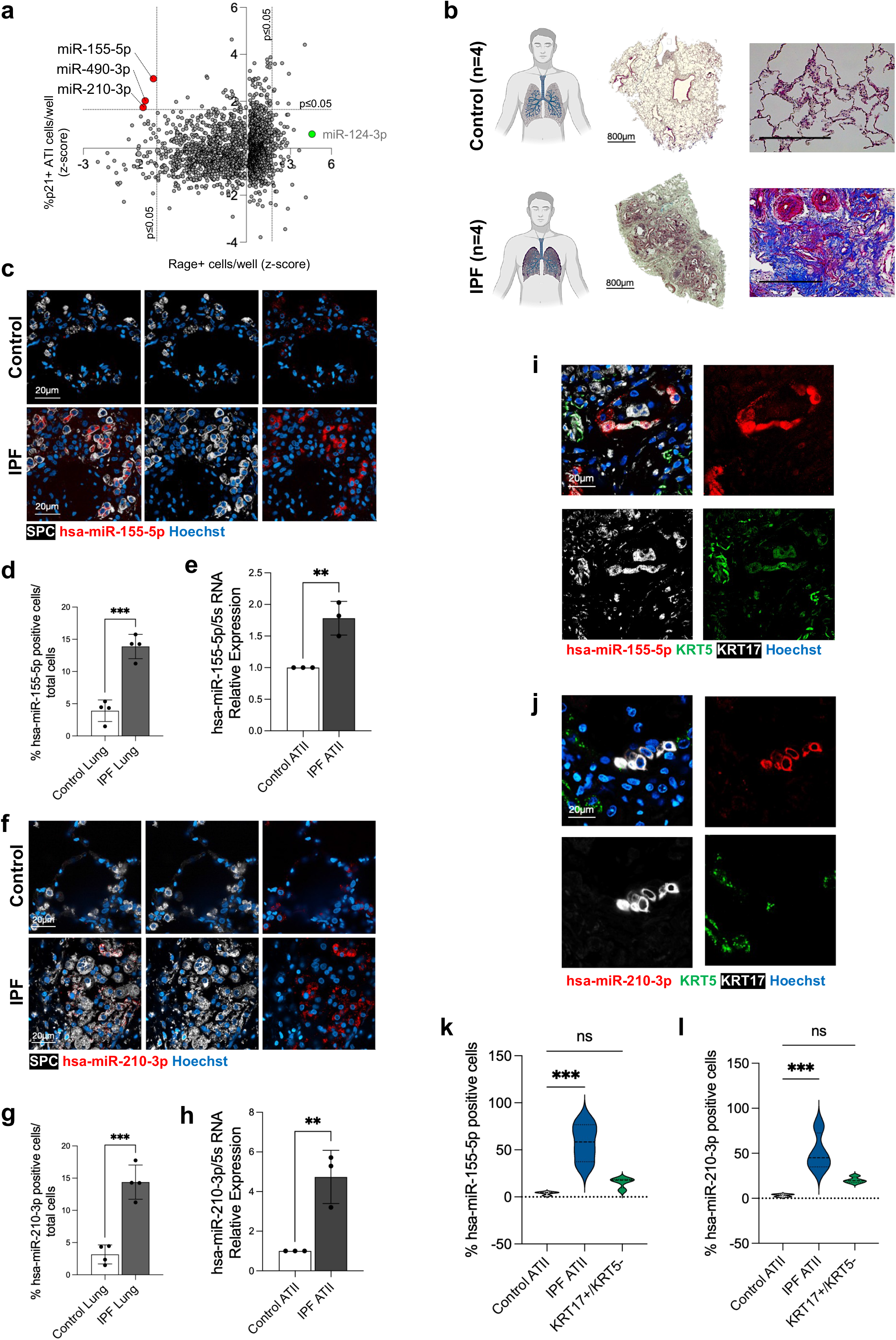
Phenotypic screening identifies miR-155-5p and miR-210-3p as disease-associated microRNAs enriched in pathological alveolar epithelial states in human IPF. a) Phenotypic screening of 2,042 human miRNA mimics in primary murine ATII cells. Candidate miRNAs were identified according to their concomitant effects on ATII-to-ATI transdifferentiation, quantified as RAGE⁺ cells per well (x axis), and cellular senescence, quantified as nuclear p21⁺ ATII cells per well (y axis). MiR-155-5p, miR-210-3p and miR-490-3p are highlighted in red, whereas miR-124-3p (control of promotion of ATII to ATI transdifferentiation) is shown in green. Dashed lines indicate the significance thresholds (P ≤ 0.05). b) Representative Masson’s trichrome staining of lung sections from healthy donors and patients with IPF (n = 4 per group), showing preserved alveolar architecture in healthy lungs and extensive fibrotic remodelling and collagen deposition in IPF lungs. Scale bars, 800 μm. c) Representative miRNA-FISH combine with immunofluorescence images of healthy and IPF human lung sections showing miR-155-5p (red), the ATII marker SPC (white) and nuclei (Hoechst, blue). Scale bars, 20 μm. d) Quantification of miR-155-5p-positive cells as a percentage of total lung cells in healthy and IPF lung tissue. e) Relative expression of mature miR-155-5p in ATII cells isolated from healthy donors and patients with IPF. f) Representative miRNA-FISH and immunofluorescence images of healthy and IPF human lung sections showing miR-210-3p (red), SPC (white) and nuclei (Hoechst, blue). Scale bars, 20 μm. g) Quantification of miR-210-3p-positive cells as a percentage of total lung cells in healthy and IPF lung tissue. h) Relative expression of mature miR-210-3p in ATII cells isolated from healthy donors and patients with IPF. i,j) Representative miRNA-FISH combine with immunofluorescence images showing miR-155-5p (i) or miR-210-3p (j) (red) within KRT17⁺/KRT5⁻ aberrant transitional epithelial cells in human IPF lungs. KRT17 is shown in white, KRT5 in green and nuclei in blue. Scale bars, 20 μm. k,l) Quantification of miR-155-5p-positive (k) and miR-210-3p-positive (l) cells across healthy ATII cells, IPF ATII cells and KRT17⁺/KRT5⁻ aberrant transitional epithelial cells. Individual data points represent independent biological samples; violin plots show the distribution of values. Statistical significance was determined using unpaired t test for d,e,g,h, and One-way ANOVA followed by Dunnett’s for k,l with multiple-comparison correction where appropriate. **P < 0.01; ***P < 0.001; ns, not significant.

We next examined miRNA expression across distinct epithelial populations in human IPF lung tissue. miR-155-5p and miR-210-3p were detected in 57.50% and 50.83% of IPF ATII cells, respectively, compared with approximately 23% of healthy ATII cells. Both miRNAs were also present in KRT17⁺/KRT5⁻ aberrant transitional epithelial cells with 15.33% ± 5.6% and 20.25% ± 3.4% of these cells expressing miR-155-5p and miR-210-3p, respectively (Fig. 1i-l).

Having established their clinical relevance, we next evaluated their therapeutic inhibition *in vitro* using ASOs in primary mouse ATII cells isolated 21 days after bleomycin injury (1U/kg). Diseased ATII cells exhibited impaired trans-differentiation and heightened senescence compared healthy ATII cells (Fig. 1a, left) and treatment with ASOs targeting either miRNA efficiently restored epithelial differentiation, evidenced by increased RAGE-positive ATI cells, while significantly reducing p21-positive senescent cells (Extended Data Fig. 2a-c). In conclusion, we identified miR-155 and miR-210 as pathogenic microRNAs that contribute to impaired ATII-to-ATI transdifferentiation and ATII cell senescence in human idiopathic pulmonary fibrosis (IPF). Furthermore, we demonstrated that in vitro inhibition of these microRNAs using antisense oligonucleotides (ASOs) is sufficient to restore efficient ATII-to-ATI transdifferentiation in diseased mouse ATII cells.

### Hypoxia induces miR-155-5p and miR-210-3p and their inhibition restores ATII-to-ATI transdifferentiation

Because miR-155-5p and miR-210-3p belong to the hypoxamiR family ^45^, we next investigated whether hypoxia directly drives their induction within the alveolar epithelium. Consistent with previous reports of dysregulated HIF signalling in experimental pulmonary fibrosis, analysis of human IPF lungs revealed predominant nuclear localization of HIF-2α and cytoplasmic accumulation of HIF-1α, a pattern consistent with HIF-2α-predominant activation reported following repetitive lung injury ^12^. (Extended Data Fig. 3a-c). However, whether hypoxia is sufficient to induce miR-155-5p and miR-210-3p specifically in primary ATII cells had not been previously demonstrated. To address this question, we isolated primary ATII cells and exposed them to hypoxic conditions (5% O₂) for 48 hours (Extended Data Fig. 3d,g). Both miRNAs were robustly induced compared with normoxic conditions (Extended Data Fig. 3e-f), establishing a direct mechanistic link between hypoxic of the fibrotic lung and the induction of these disease-associated microRNAs within the alveolar epithelium.

We next investigated the functional consequences of this previously unexplored epithelial HIF–hypoxamiR axis. ATII cells were transfected with ASOs targeting either miRNA before hypoxic exposure and subsequently exposed to hypoxia (5% O₂) (Extended Data Fig. 3h). ASO treatment efficiently reduced the expression of both miRNAs (Extended Data Fig. 3i-j) and significantly decreased HIF-1α expression (Extended Data Fig. 3k), indicating disruption of a HIF-dependent feed-forward regulatory loop. Functionally, inhibition of either miRNA rescued ATII cell survival, restored total cell numbers, and re-established ATII-to-ATI transdifferentiation under hypoxic conditions (Extended Data Fig. 3l-n).

### miR-155-5p and miR-210-3p mirror fibrosis progression and age-dependent lung regenerative failure

We next investigated whether the expression of miR-155-5p and miR-210-3p correlates with disease progression in vivo. To track miRNA dynamics across the full fibrosis-repair cycle, we further quantified their expression in young mice treated with bleomycin (Extended Data Fig. 4). Histological and biochemical analysis of lung tissue collected between days 10 and 60 confirmed progressive fibrotic remodeling, which peaked between days 21 and 30 before resolving during the repair phase (days 45– 60) (Extended Data Fig. 4b-c). Notably, the expression of both miRNAs closely mirrors disease progression, increasing during fibrosis and declining during resolution (Extended Data Fig. 4d-e). Notably, HIF-1α induction preceded hypoxamiR expression, peaking at day 10 (Extended Data Fig. 4 g, j, m), whereas miR-155-5p and miR-210-3p reached maximal levels between days 21 and 30 (Extended Data Fig. 4d-e), coincident with peak fibrotic remodelling. This temporal sequence suggests that an early hypoxic response drives subsequent hypoxamiR induction, thereby sustaining a dysfunctional ATII cell state characterized by induction of the senescent markers p21 and p16 (Extended Data Fig. 4 f, i, l).

Aged mice, unlike young mice, which typically develop transient, self-resolving fibrosis following bleomycin injury, fail to regenerate alveolar structures after bleomycin treatment usually displaying more severe lung damage than young animals ^46–48^ (Extended Data Fig. 5a-c). Accordingly, aged mice exhibited elevated basal expression of both miR-155-5p and miR-210-3p, which increased further following bleomycin injury (Extended Data Fig. 5d-e).

Collectively, these findings identify miR-155-5p and miR-210-3p as key mediators of alveolar epithelial dysfunction and provide a strong rationale for evaluating whether their therapeutic inhibition can promote lung repair in vivo.

### Inhibition of miR-155-5p and miR-210-3p reverses established fibrosis and restores alveolar epithelial homeostasis in aged mice

Having identified miR-155-5p and miR-210-3p as clinically relevant regulators of pathological epithelial states, we next asked whether their therapeutic inhibition could reverse established fibrosis and restore alveolar homeostasis in vivo. Because IPF predominantly affects elderly individuals, we used 20-month-old mice, a clinically relevant model characterized by persistent, non-resolving fibrosis and defective alveolar regeneration ^46^. Aged mice received intratracheal ASOs targeting miR-155-5p or miR-210-3p beginning ten days after bleomycin injury, when fibrosis was already established (Fig. 2a). By day 21, vehicle-treated mice developed extensive architectural remodelling and collagen deposition, whereas both ASOs markedly reduced fibrosis, lung weight and hydroxyproline content (Fig. 2b–e). Hydroxyproline levels strongly correlated with miR-155-5p and miR-210-3p expression (Fig. 2d,e), directly linking hypoxamiR abundance to fibrotic burden. Unexpectedly, inhibition of either miRNA reduced both hypoxamiRs (Fig. 2d,e), suggesting a shared regulatory circuit, although the two ASOs produced distinct architectural outcomes: miR-155-5p inhibition preferentially restored compact alveolar architecture despite residual interstitial fibrosis, whereas miR-210-3p inhibition resulted in minimal fibrosis and larger alveolar spaces.

**Figure 2.**
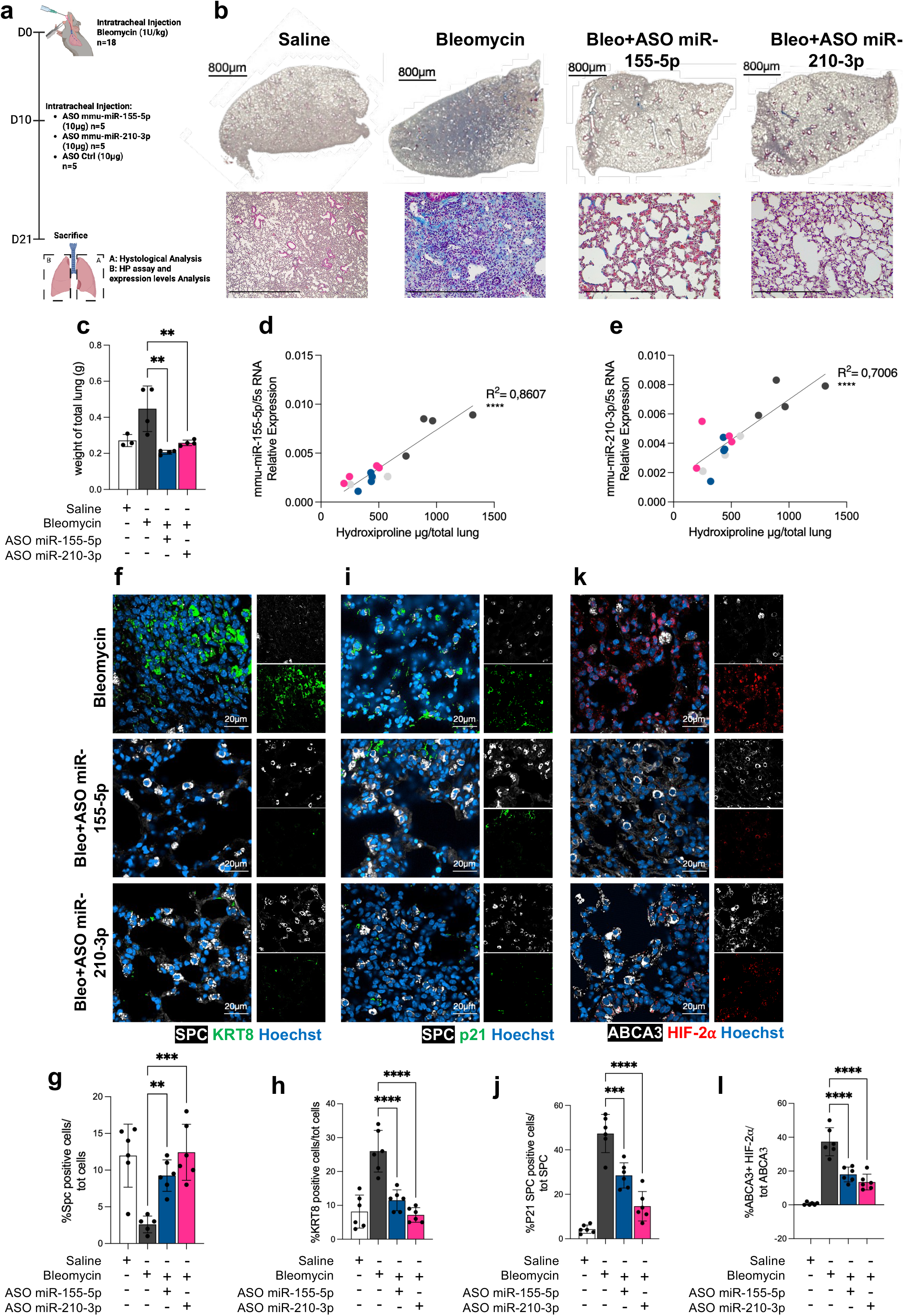
Therapeutic inhibition of miR-155-5p and miR-210-3p reverses established pulmonary fibrosis and restores alveolar epithelial homeostasis in aged mice 20 months. a) Schematic representation of the therapeutic protocol. Aged mice received intratracheal bleomycin (1 U kg⁻¹) on day 0, followed by intratracheal administration of ASOs targeting miR-155-5p or miR-210-3p (10 μg) on day 10; lungs were collected on day 21 for histological, biochemical and molecular analyses. b) Representative Masson’s trichrome staining of lungs from saline, bleomycin, bleomycin + ASO-miR-155-5p and bleomycin + ASO-miR-210-3p treated mice. Whole-lung sections and higher-magnification regions are shown. Scale bars, 800 μm for whole-lung sections. c) Lung weight across the indicated experimental groups. d,e) Correlation between lung hydroxyproline content and miR-155-5p (d) or miR-210-3p **(e)** expression across experimental conditions. Colours denote the corresponding treatment groups. R² values are indicated. **f)** Representative immunofluorescence images showing SPC (white), KRT8 (green) and nuclei (Hoechst, blue) in bleomycin- and ASO-treated lungs. Scale bars, 20 μm. **g)** Quantification of SPC⁺ cells as a percentage of total cells. **h)** Quantification of KRT8⁺ cells as a percentage of total cells. **i)** Representative immunofluorescence images showing SPC (white), p21 (green) and nuclei (Hoechst, blue). **j)** Quantification of p21⁺/SPC⁺ cells relative to the total SPC⁺ population. **k)** Representative immunofluorescence images showing the ATII marker ABCA3 (white), HIF-2α (red) and nuclei (Hoechst, blue). **l)** Quantification of ABCA3⁺/HIF-2α⁺ cells relative to the total ABCA3⁺ population. Individual points represent biological replicates. Statistical significance was determined using One-way ANOVA followed by Dunnett’s for c, g, h, j, l with multiple-comparison correction where appropriate. **P < 0.01; ***P < 0.001; ****P < 0.0001.

We next determined whether fibrosis reversal was accompanied by restoration of epithelial homeostasis. Bleomycin induced a marked accumulation of KRT8⁺ transitional epithelial cells, the murine counterpart of the KRT17⁺/KRT5⁻ aberrant epithelial intermediates found in human IPF ^6^, whereas either ASO reduced KRT8⁺ cell abundance and Krt8 expression while increasing the SPC⁺ ATII population (Fig. 2f–h). HypoxamiR inhibition also reduced p21⁺ epithelial cells and attenuated p16 expression across experimental models (Fig. 2i,j and Extended Data Fig. 5g), consistent with resolution of epithelial senescence. Finally, ASO treatment reduced HIF1A expression (Extended Data Fig. 5h) and nuclear HIF-2α accumulation in SPC⁺ ATII cells (Fig. 2k,l), indicating normalization of pathological HIF signalling.

To complement these efficacy studies with an assessment of therapeutic safety and pulmonary delivery, we evaluated ASO tolerability and biodistribution in healthy cells and young mice. ASO treatment did not alter the morphology, differentiation or phenotype of healthy primary ATII cells or lung fibroblasts (Extended Data Fig. 6a–f), nor did it increase proliferation of epithelial-like lung adenocarcinoma cells (A549) expressing basal levels of both hypoxamiRs (Extended Data Fig. 6g,h). Following pulmonary administration in young mice, ASOs efficiently reached the alveolar epithelium, with 22.10 ± 0.14% of ATII cells labelled by flow cytometry and 34.39 ± 6.7% by immunofluorescence (Extended Data Fig. 7a–f), and remained detectable in the lung for up to 60 days after a single administration without detectable toxicity in extrapulmonary organs (Extended Data Fig. 7i–l). Together, these findings support sustained pulmonary delivery and tolerability of hypoxamiR-targeting ASOs.

Although hypoxamiR inhibition reversed established fibrosis and restored epithelial homeostasis, the cellular mechanisms driving this therapeutic response remained unclear. Specifically, whether ASO treatment enables dysfunctional ATII cells to re-enter regenerative programmes and rebuild alveolar structures, and how this epithelial response contributes to the removal of established fibrosis, remained unknown. We therefore sought to directly investigate the cellular dynamics underlying ASO-mediated alveolar repair and remodelling of established fibrosis.

### Live imaging of lineage traced ATII cells reveals rapid reactivation of alveolar epithelial regeneration following miRNA inhibition

Previous experiments demonstrated that ASO-mediated inhibition of miR-155-5p and miR-210-3p not only reversed established fibrosis but also restored the ATII compartment and alveolar architecture (Fig. 2; Extended Data Fig. 5). We therefore sought to directly visualize the early epithelial dynamics underlying alveolar repair.

To this end, we combined genetic lineage tracing with live imaging of precision-cut lung slices (PCLS) from Sftpc-CreERT2; Rosa26-mTmG mice, enabling longitudinal tracking of individual GFP-labelled ATII cells during regeneration rather than static endpoint analysis. Mice received tamoxifen followed by a 10-day washout, bleomycin injury and intratracheal ASO administration 10 days later. Five days after treatment, lungs were collected and PCLS were subjected to live imaging (Fig. 3a, b). Quantitative single-cell tracking revealed distinct epithelial responses to ASO miR-155-5p and ASO miR-210-3p inhibition (Fig. 3c–f). MiR-155-5p inhibition preferentially promoted morphological remodeling, with GFP⁺ cells displaying increased elongation (c), cell area (d) total distance travelled (f), and migration speed, whereas miR-210-3p inhibition produced a similar but different motility phenotype, characterized by increased cell area (d) as well as total distance travelled (e), but reduced migration speed. These dynamic differences were consistent with the time-lapse imaging, indicating that the two ASOs engage distinct but complementary regenerative behaviours (Fig. 3b–f).

**Figure 3.**
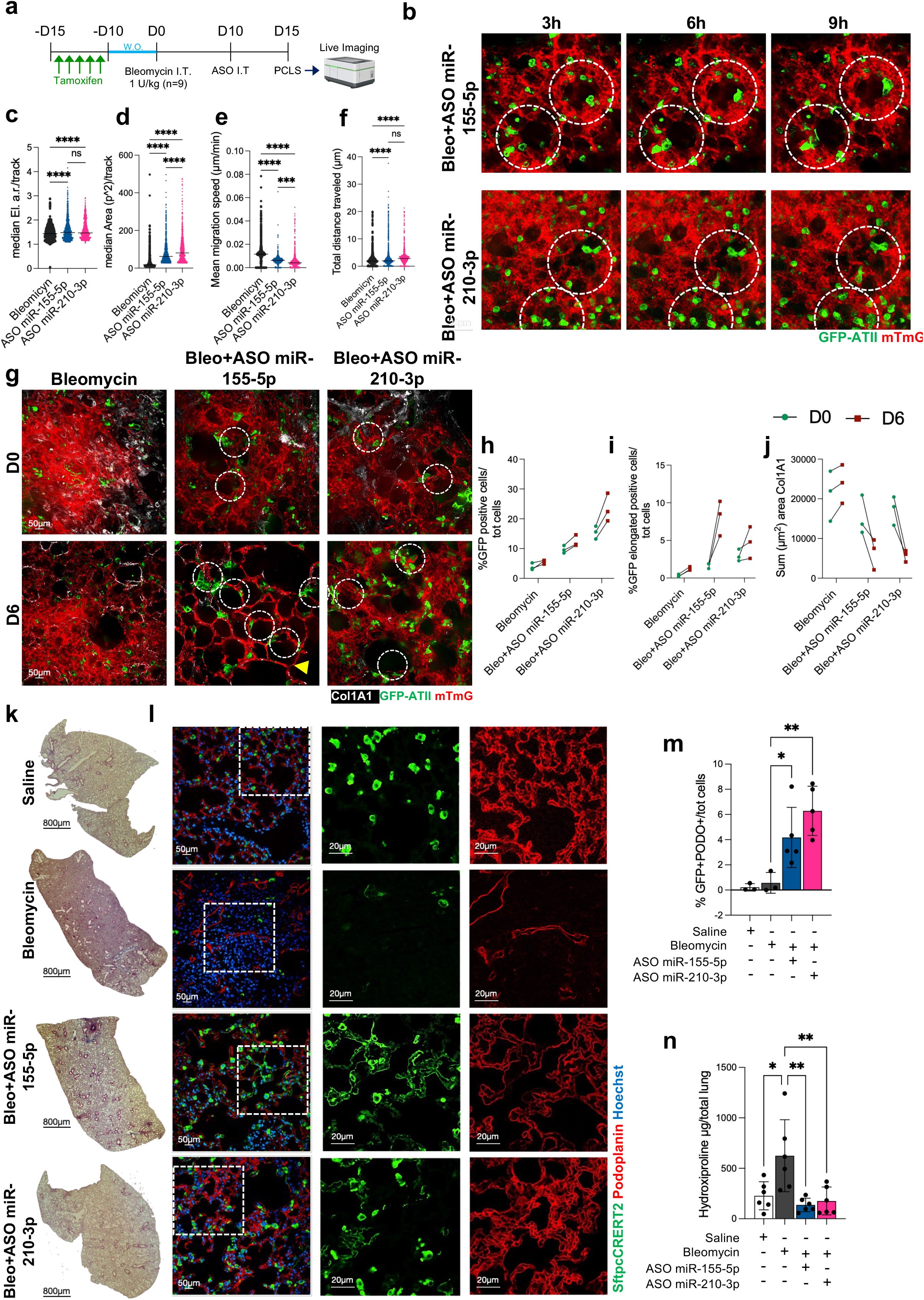
Live PCLS imaging and lineage tracing reveal restoration of ATII regenerative dynamics and ATII-to-ATI differentiation following hypoxiamiR inhibition. a) Experimental design for lineage tracing and live imaging of ATII cells in precision-cut lung slices (PCLS) from *Sftpc-CreERT2;Rosa26-mTmG* mice following bleomycin injury and intratracheal administration of ASO miR-155-5p or ASO miR-210-3p after 10 days from Bleomycin injection. Mice were sacrifice after 5 days from ASOs treatment b) Representative time-lapse images (after 3h, 6h and 9h of acquisition) of GFP-labelled ATII cells in PCLS following ASO miR-155-5p or ASO miR-210-3p treatment. Dashed circles highlight representative GFP⁺ cells undergoing morphological changes or displacement over time. c–f) TrackMate-based single-cell analysis of GFP⁺ ATII cells showing median ellipse aspect ratio (c), median cell area (d), mean migration speed (e), and total distance travelled (f) in bleomycin-, ASO miR-155-5p- and ASO miR-210-3p-treated PCLS. Each point represents an individual tracked cell; distributions are shown as violin plots. g) Representative fluorescence images of PCLS at D0 and D6 showing lineage-labelled GFP⁺ ATII cells (green), Col1a1 (white) and mTmG-labelled tissue (red); dashed circles indicate representative GFP⁺ epithelial cells. h–j) Quantification of GFP⁺ cells as a percentage of total cells (h), elongated GFP⁺ cells as a percentage of total cells (i), and Col1a1-positive area (**j**) at D0 and D6. **k)** Representative whole-lung histological sections from saline-, bleomycin-, ASO miR-155-5p- and ASO miR-210-3p-treated mice. **l)** Representative immunofluorescence images showing Sftpc (green), PDPN/podoplanin (red) and Hoechst (blue), with enlarged views illustrating epithelial remodeling. **m)**Quantification of GFP⁺PDPN⁺ cells as a percentage of total cells. **n)** Lung hydroxyproline content across treatment groups. Statistical significance is indicated in the figure; ns, not significant; *P < 0.05, **P < 0.01, ****P < 0.0001.

Endpoint analysis supported these divergent trajectories: both ASOs treatments increased the GFP⁺ epithelial compartment, while miR-155-5p inhibition produced the strongest increase in elongated GFP⁺ cells (Fig. 3g–i). Both treatments were accompanied by marked remodelling of the fibrotic niche, including reduced Col1a1-positive matrix deposition (Fig. 3g, j). Together, these findings suggest that miR-210-3p ASO significantly increased GFP⁺ cell numbers (Fig. 3c, d), while miR-155-5p inhibition predominantly promotes epithelial morphological maturation, whereas miR-210-3p inhibition favours a more migratory regenerative response, with both programs ultimately contributing to alveolar repair.

### Inhibition of miR-155-5p and miR-210-3p restores ATII-to-ATI differentiation and regenerates alveolar architecture in vivo

Live imaging established that hypoxamiR inhibition rapidly reactivated epithelial dynamics. We next asked whether these early cellular responses ultimately generated mature alveolar structures in vivo. Using the same ATII lineage tracing model, ASOs were administered intratracheally 21 days after injury, when fibrosis was fully established (Extended Data Fig 4a, b), and lungs were analysed nine days later (Fig. 3h).

Consistent with our previous findings, ASO treatment significantly reduced fibrotic burden, as demonstrated by decreased hydroxyproline content and collagen deposition (Fig. 3g, j). More importantly, lineage tracing demonstrated that epithelial regeneration was strongly promoted by the treatment. Immunofluorescence analysis revealed a marked increase in GFP⁺/PDPN⁺ double-positive cells following treatment (Fig. 3h, i), indicating successful differentiation of lineage-labelled ATII cells into mature ATI cells. This regenerative response was accompanied by a significant reduction in persistent KRT8⁺ transitional epithelial cells, indicating resolution of the pathological epithelial intermediate state that characterizes fibrotic lungs.

The simultaneous expansion of ATII-derived mature ATI cells together with loss of pathological KRT8-positive intermediates demonstrates that selected miRNA inhibition restores ATII plasticity, allowing pathological epithelial cells to complete terminal differentiation and rebuild functional alveolar architecture. Thus, ASO therapy does not simply halt fibrosis progression but actively reawakens the endogenous regenerative programme of the adult lung.

### Hypoxia-induced miR-155-5p and miR-210-3p couple epithelial dysfunction to fibroblast activation through paracrine signalling

As previously shown, we demonstrated that both miR-155-5p and miR-210-3p contribute directly to fibroblast activation. Lung fibroblasts isolated from bleomycin-treated mice exhibited increased expression of both miRNAs (Extended Data Fig. 1g,k-l), while forced overexpression of either miRNA promoted myofibroblast differentiation and cellular senescence (Extended Data Fig. 1m-o). Conversely, ASO treatment attenuated TGF-β-induced myofibroblast differentiation in primary lung fibroblast (Extended Data Fig. 2d-j).

However, following intratracheal ASO delivery, approximately 34% of ATII cells internalized ASOs, whereas only 11.2% of lung fibroblasts were targeted (Extended Data Fig. 7d–f), a level unlikely to account for the profound reduction in fibrosis through direct fibroblast reprogramming alone.

Therefore, we hypothesized that pathological ATII cells overexpressing both miR-155-5p and miR-210-3p are not only impaired in their regenerative capacity but also actively promote profibrotic remodelling by driving fibroblast activation through paracrine signalling.

To directly test this possibility, we investigated whether the secretome of miRNA-reprogrammed ATII cells was sufficient to drive fibroblast activation in the absence of direct cell-to-cell contact (Fig. 4a). Conditioned medium from ATII cells overexpressing the selected miRNAs robustly induced fibroblast-to-myofibroblast activation, as demonstrated by the acquisition of a typical myofibroblast morphology, increased α-SMA expression, and enhanced cellular senescence, as indicated by p21 upregulation (Fig. 4b-d). To investigate the nature of the mediators responsible for this effect, the conditioned medium was fractionated into three molecular weight ranges: proteins (50–10 kDa), peptides (10–3 kDa), and low-molecular-weight metabolites (<3 kDa).

**Figure 4.**
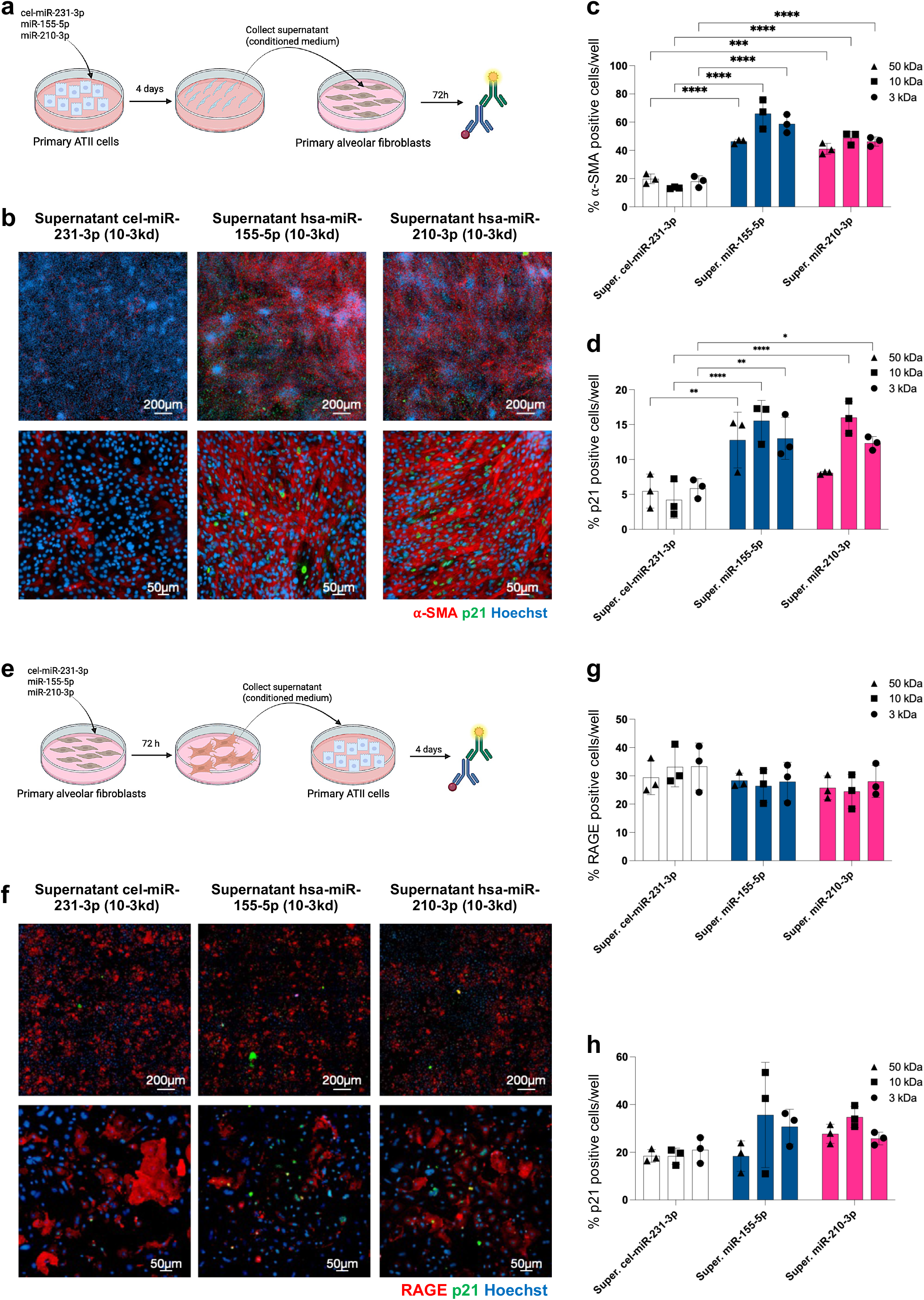
HypoxamiR-dependent bidirectional paracrine crosstalk between alveolar epithelial cells and fibroblasts. **a)** Schematic representation of the ATII-to-fibroblast conditioned-medium transfer experiment. Primary murine ATII cells were transfected with cel-miR-231-3p, hsa-miR-155-5p or hsa-miR-210-3p. After 4 days, conditioned medium was collected, fractionated using the indicated molecular-weight cut-offs and transferred to primary alveolar fibroblasts for 72 h before fixation and analysis. **b)** Representative immunofluorescence images of primary alveolar fibroblasts exposed to the 10-kDa conditioned-medium fraction from control, miR-155-5p or miR-210-3p transfected ATII cells and stained for α-SMA (red), p21 (green) and Hoechst (blue). **c,d)** Quantification of α-SMA-positive (**c**) and p21-positive (**d**) fibroblasts following exposure to the 50-, 10- and 3-kDa conditioned-medium fractions from the indicated ATII-cell conditions. **e)** Schematic representation of the reciprocal fibroblast-to-ATII conditioned-medium transfer experiment. Primary alveolar fibroblasts were transfected with cel-miR-231-3p, hsa-miR-155-5p or hsa-miR-210-3p. Conditioned medium was collected after 72 h, fractionated using the indicated molecular-weight cut-offs and transferred to primary ATII cells for 4 days before fixation and analysis. **f)** Representative immunofluorescence images of primary ATII cells exposed to the 10-kDa conditioned-medium fraction from control-, miR-155-5p- or miR-210-3p-transfected fibroblasts and stained for RAGE (red), p21 (green) and Hoechst (blue). **g,h)** Quantification of RAGE-positive (**g**) and p21-positive (**h**) ATII cells following exposure to the 50-, 10- and 3-kDa conditioned-medium fractions from the indicated fibroblast conditions. Scale bars, 200 μm (upper images in **b,f**) and 50 μm (lower images in **b,f**). Statistical significance was determined using one-way ANOVA followed by Tukey’s multiple comparisons test for c,d,g,h with multiple-comparison correction where appropriate. *P < 0.05; **P < 0.01; ***P < 0.001; ****P < 0.0001.

Remarkably, the biological activity was preserved across all fractions and was maintained, or even enhanced, in the lower molecular weight fractions. These findings indicate that fibroblast activation is mediated predominantly by low-molecular-weight soluble factors rather than by large secreted proteins (Fig. 4c-d).

We next examined the reciprocal paracrine interaction by exposing primary mouse ATII cells to conditioned medium from alveolar fibroblasts overexpressing the selected miRNAs (Fig. 4e). In contrast to the robust epithelial-to-mesenchymal signalling observed above, conditioned medium from miRNA-overexpressing fibroblasts induced only a modest, non-significant reduction in ATII-to-ATI transdifferentiation and a slight, non-significant increase in p21 positivity (Fig. 4f–h).

Given the central role of miRNA-overexpressing ATII cells in remodelling the fibrotic microenvironment, it remained unclear how ASO-treated ATII cells exit this altered state and regain their capacity to support effective alveolar repair. We therefore sought to define the transcriptional network restored by hypoxamiR inhibition in vivo.

### ASO therapy rewires the ATII transcriptional programme to restore alveolar regeneration

Given the pleiotropic nature of miRNA regulation, we next defined the molecular programmes underlying epithelial reprogramming following hypoxamiR inhibition. Lineage-traced ATII cells were isolated from bleomycin-treated Sftpc-mTmG mice following therapeutic administration of ASO-miR-155-5p or ASO-miR-210-3p (Fig. 5a).

**Figure 5.**
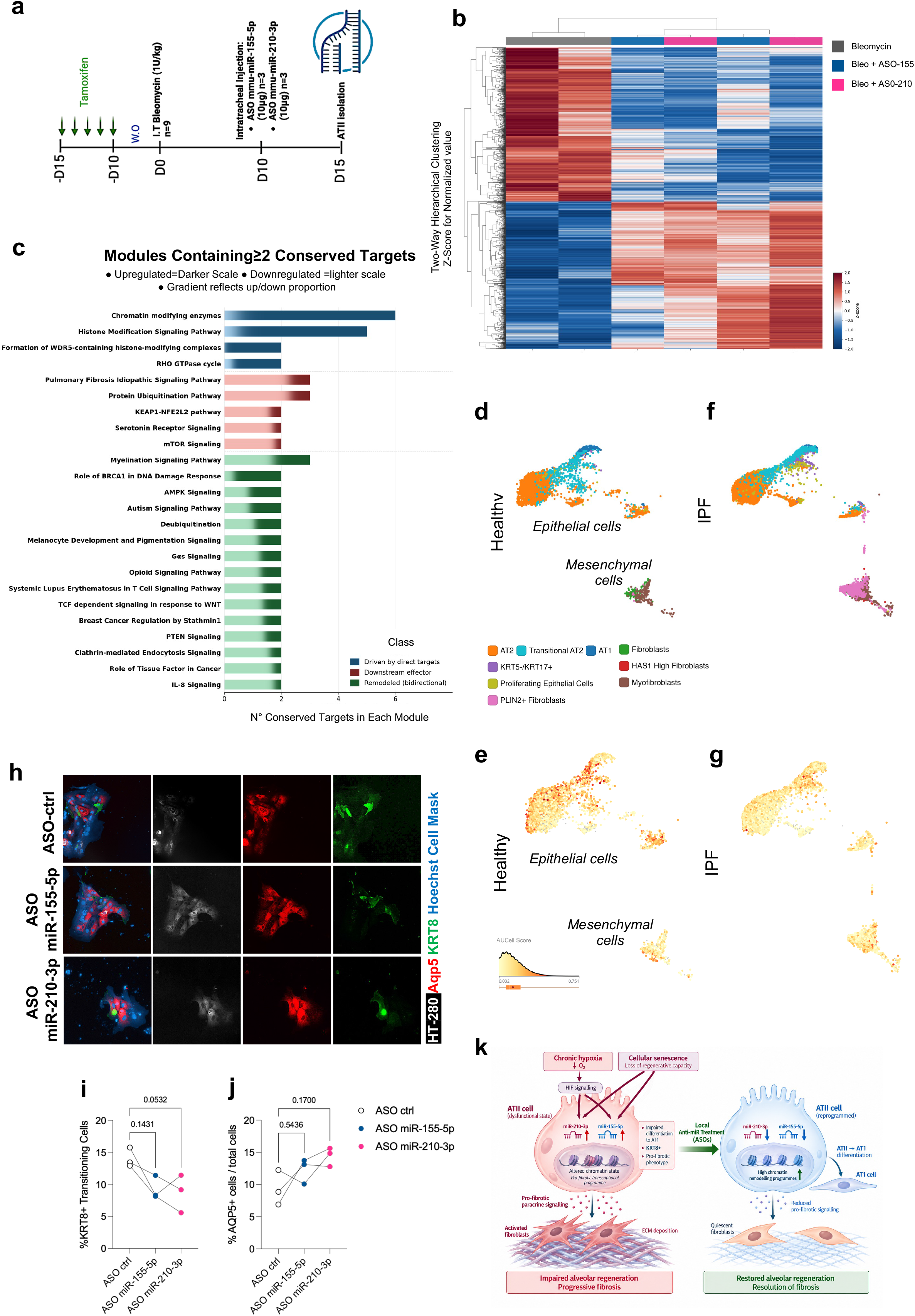
HypoxamiR inhibition restores a conserved alveolar epithelial programme disrupted in human IPF. **a)** Experimental design for transcriptomic profiling of lineage-traced alveolar type II (ATII) cells. *Sftpc-CreERT2;Rosa-mTmG* mice received tamoxifen before intratracheal bleomycin administration (1 U kg−1), followed by therapeutic intratracheal administration of ASO-miR-155-5p or ASO-miR-210-3p (10 μg per mouse) at day 10. Lineage-labelled ATII cells were isolated at day 15 for transcriptomic analysis. **b)** Unsupervised hierarchical clustering of bulk RNA-sequencing profiles from lineage-traced ATII cells isolated from bleomycin-treated mice and mice treated with ASO-miR-155-5p or ASO-miR-210-3p. Heatmap shows row-normalized *z*-scores for 1977 differentially expressed genes. **c)** Canonical pathways associated with the transcriptional response to hypoxamiR inhibition and their representation by conserved candidate miRNA targets. Bars indicate the number of genes within each pathway that were commonly upregulated (dark shading) or downregulated (light shading) following ASO treatment. Colour intensity reflects gene counts, with darker shades indicating higher counts, lighter shades indicating lower counts. Pathways are grouped according to their predominant relationship to direct miRNA regulation and broader transcriptional remodelling. Downstream effector in red, Driven by direct targets in blue and Remodeled in green **d,f)** UMAP representations of selected epithelial and mesenchymal populations from healthy (**d**) and IPF (**f**) human lungs in the Habermann *et al.* scRNA-seq dataset (GSE135893), coloured according the legend. **e,g)** Distribution of the ASO-restored hypoxamiR target-gene programme across the corresponding healthy (**e**) and IPF (**g**) epithelial and mesenchymal compartments. **h)** Representative immunofluorescence images of primary ATII cells isolated from n=3 patients with IPF and treated with control ASO, ASO-miR-155-5p or ASO-miR-210-3p. Cells were stained for HTII-280 (white), AQP5 (red), KRT8 (green) and nuclei (Hoechst, blue); the merged image is shown on the left. **i)** Quantification of HTII-280+ AQP5+ KRT8+ cells following ASO treatment. **j)** Quantification of %AQP5+ cells following ASO treatment. *n* = 3 independent human IPF samples. Data are mean ± s.d.; individual points represent biological replicates. **k)** Working model of hypoxia– hypoxamiR signalling and its therapeutic inhibition. Chronic hypoxia induces miR-155-5p and miR-210-3p, promoting ATII regenerative failure, accumulation of pathological transitional epithelial states, senescence and profibrotic epithelial–mesenchymal signalling. Therapeutic hypoxamiR inhibition restores target-gene expression and epithelial plasticity, favouring ATII-to-ATI differentiation and alveolar regeneration. Statistical significance was assessed by one-way-repeated-measures ANOVA followed by Dunnett’s multiple-comparisons test. *P* values are reported.

RT–qPCR confirmed target engagement (Extended Data Fig. 8a,b), and bulk mRNA sequencing identified differentially expressed protein-coding genes. Both ASOs induced highly convergent transcriptional responses that segregated from bleomycin controls, consistent with restoration of a common epithelial state (Fig. 5b).

To distinguish programmes directly released by miRNA inhibition from secondary transcriptional responses, we integrated genes commonly regulated by both ASOs with conserved miR-155-5p and miR-210-3p candidate direct targets derepressed following treatment. Among 24 pathways meeting these criteria, chromatin-modifying enzymes and histone modification signalling ranked highest, with broad pathway upregulation and six conserved candidate targets spanning histone methylation, demethylation, acetylation and chromatin remodelling (Fig. 5c and Supplementary Table 1). This hierarchy was preserved using human-only or mouse-only target predictions (Spearman ρ = 0.624–0.695, all P < 0.001; Kendall’s W = 0.553, P = 0.024; Extended Data Fig. 8c,d). By contrast, IPF signalling was predominantly characterized by downregulated transcripts and contained few candidate miRNA targets, consistent with secondary normalization following epithelial reprogramming (Fig. 5c).

We next assessed conservation of the ASO-restored programme across independent murine fibrosis and human IPF single-cell RNA-sequencing datasets ^4,6–8,44,49^. Across datasets, transcripts induced by ASO treatment were reciprocally downregulated in fibrotic ATII cells, with ∼45–72% concordance across human IPF datasets and ∼83– 90% across murine fibrosis datasets for both miR-155-5p-and miR-210-3p-regulated genes (Extended Data Fig. 8e–g). In human IPF lungs, expression of 30 selected common candidate targets was preferentially reduced in diseased epithelial populations while remaining comparatively preserved in fibroblasts (Fig. 5d–g), linking the ASO-restored programme to the epithelial compartment disrupted in IPF.

We therefore asked whether targeting these hypoxamiRs could functionally reverse the aberrant epithelial phenotype directly in primary cells from patients with IPF. Consistent with persistent transitional epithelial states in human IPF, primary IPF ATII cultures contained KRT8-positive transitional cells ^7,50^. Treatment with either ASO-miR-155-5p or ASO-miR-210-3p reduced the KRT8-positive transitional population, accompanied by an increase in AQP5-positive cells, indicative of acquisition of an ATI phenotype (Fig. 5h–j). Thus, the transcriptional programme identified across mouse and human datasets translated into phenotypic rescue in patient-derived alveolar epithelial cells, supporting the ability of hypoxamiR inhibition to release a disease-associated transitional state and promote alveolar differentiation.

Together, these findings support a model in which chronic hypoxia-induced miR-155-5p and miR-210-3p constrain ATII regenerative competence through a transcriptional programme converging on chromatin regulation. Therapeutic hypoxamiR inhibition releases this programme, restores epithelial plasticity and redirects the fibrotic response towards alveolar regeneration (Fig. 5k).

## Discussion

Our findings identify miR-155-5p and miR-210-3p as therapeutically accessible targets for the treatment of IPF, linking impaired alveolar epithelial regeneration to profibrotic signalling. The strength of this conclusion lies in the convergence of functional discovery, therapeutic intervention in preclinical models and relevance to the human disease.

Phenotype-guided re-analysis of a previously reported functional screen of 2,042 human miRNA mimics in primary murine ATII cells^32^ prioritized miR-155-5p and miR-210-3p as regulators that impaired ATII-to-ATI differentiation while increasing p21 positivity, thereby promoting an aberrant transitional state characterized by regenerative arrest, senescence and profibrotic paracrine signalling. In this model, dysfunctional ATII cells not only fail to progress along the ATII-to-ATI regenerative trajectory but also sustain pathological epithelial–mesenchymal crosstalk that promotes fibroblast activation.

Therapeutic inhibition of miR-155-5p or miR-210-3p disrupted both components of this response, restoring ATII-derived alveolar regeneration while reducing fibrosis in vivo. Notably, these effects were maintained in aged mice, a setting in which the regenerative response to lung injury is intrinsically impaired and that therefore more closely reflects the age-associated loss of epithelial resilience characteristic of IPF ^46-48^.

The relevance of this mechanism to human disease is particularly important given the increasing recognition of IPF as a disease of failed ATII-driven alveolar regeneration rather than primarily fibroblast-driven fibrosis ^51,52^. Both miR-155-5p and miR-210-3p were increased in failing ATII cells from patients with IPF and enriched in KRT17⁺/KRT5⁻ aberrant epithelial cells. Conversely, the transcriptional programme restored by ASO treatment was consistently suppressed in fibrotic ATII cells across independent human IPF datasets, and direct inhibition of either miRNA in primary ATII cells from patients with IPF reduced aberrant KRT8-positive transitional cells and promoted acquisition of an ATI phenotype. Nevertheless, cultured cells capture only part of the diseased alveolar environment, and the limited number of donors and heterogeneity of IPF warrant broader assessment to establish the consistency of this response across patients, disease stages and treatment backgrounds. Together, these observations link hypoxia-induced miR-155-5p and miR-210-3p expression to a dysfunctional ATII cell state in human disease.

Anti-miRNA-ASOs, unlike replacement strategies requiring delivery of an exogenous miRNA, suppress pathologically elevated endogenous transcripts and can be chemically optimized for stability and tissue delivery. Oligonucleotide inhibition of miRNAs has already proved clinically feasible, including miravirsen-mediated inhibition of miR-122 ^42^, and repeated anti-miR dosing has been evaluated safe in humans in other disease settings ^28,43^. This distinction may be relevant given the safety limitations encountered with miRNA-replacement approaches ^53,54^. Together with the therapeutic efficacy achieved through local pulmonary delivery in our models, these properties support the translational potential of anti-miRNA ASOs to address an unmet therapeutic need in IPF by targeting epithelial dysfunction and regenerative failure, which are not directly addressed by current approved therapies, including pirfenidone, nintedanib and nerandomilast ^15,55^.

Mechanistically, microRNAs act through distributed target networks rather than through a single dominant effector, making attribution of the overall phenotype to an individual target challenging. This principle is shared by other pleiotropic therapeutics, including glucocorticoids, which exert broad transcriptional effects and remain clinically effective, and widely used, despite incompletely resolved molecular mechanisms of action ^56,57^.

In this context, our transcriptomic analyses provide a starting point for mechanistic dissection of the epithelial response to miR-155-5p and miR-210-3p inhibition. Integration of transcriptomic and target-prediction analyses revealed convergence of miR-155-5p and miR-210-3p regulation on chromatin-associated programmes supporting broader induction of both permissive and repressive chromatin regulators. Rather than indicating a uniformly permissive epigenetic state, these findings suggest that hypoxamiR inhibition may restore the chromatin-regulatory capacity required for ATII plasticity. By contrast, fibrosis-associated signalling pathways contained comparatively few candidate miRNA targets despite extensive transcriptional normalization, consistent with their regulation downstream of this broader epithelial reprogramming.

While our findings provide robust and novel insights into the role of hypoxia-induced miR-155-5p and miR-210-3p in promoting ATII cell dysfunction in IPF, several important questions remain unresolved. Establishing whether chromatin remodelling is required for epithelial rescue will require direct profiling of chromatin accessibility and histone states, together with perturbation of individual or cooperative candidate regulators. The epithelial factors mediating the profibrotic activity of the pathological ATII secretome also remain to be defined. Finally, although histological, molecular and lineage-tracing analyses demonstrated structural and regenerative rescue, pulmonary function was not assessed, and the contribution of other components of the alveolar niche, particularly the vasculature, remains unexplored.

Collectively, our findings identify miR-155-5p and miR-210-3p as key effectors of chronic hypoxic signalling that lock ATII cells into a dysfunctional state characterized by a dual pathological phenotype: impaired capacity to support effective alveolar regeneration and sustained pro-fibrotic remodelling through paracrine signalling. Therapeutic hypoxamiR inhibition releases ATII cells from this maladaptive state, restores epithelial plasticity, and shifts the injured alveolar niche from fibrosis towards regeneration, providing a mechanistic rationale for epithelial-directed regenerative therapy in IPF.

## Methods

### Study design

The study combined phenotypic re-analysis of a pre-existing miRNA screening dataset, primary-cell assays, bleomycin-induced pulmonary fibrosis models, lineage tracing and precision-cut lung slice (PCLS) imaging, human IPF tissue and primary-cell validation, and bulk and single-cell transcriptomic analyses to define the role of miR-155-5p and miR-210-3p in alveolar epithelial regenerative failure and therapeutic response. Experimental group sizes, independent biological replicates and statistical tests are reported in the figure legends and Source Data. No data points were excluded unless specified a priori by assay-quality criteria.

### Animal studies

All animal experiments were conducted in compliance with the European guidelines and international laws and policies, with the approval of ICGEB Animal Welfare Board, Ethical Committee, and Italian Ministry of Health (Authorization n° 1053/2024-PR (prot. D441B.66) and Authorization n°811/2025-PR (prot. D441B.72).

#### Bleomycin-induced pulmonary fibrosis

Male C57BL/6, Sftpc-CreERT2 and Rosa-mTmG mice aged 8 weeks or 20 months were maintained under controlled environmental conditions on a 12-h light/dark cycle. Mice were anaesthetized with ketamine (80–100 mg kg−1) and domitor (0.8–1 mg kg−1) and received bleomycin intratracheally in 60 µl. Young mice received 1 U kg−1 bleomycin and aged mice received 0.8 U kg−1. Anaesthesia was reversed with atipamezole (5 mg kg−1). Animals were monitored throughout the study and euthanized at the indicated time points. At collection, lungs were perfused with PBS; tissue was allocated to histology, immunofluorescence, RNA analysis, hydroxyproline quantification, flow cytometry or primary-cell isolation as required.

#### In vivo ASO administration and ATII lineage tracing

Antisense oligonucleotide inhibitors targeting miR-155-5p or miR-210-3p were administered intratracheally at 10 µg per mouse at day 10 or day 21 after bleomycin injury, according to the experimental design. Mice were analysed at day 21 or day 30, respectively. For lineage tracing, Sftpc-CreERT2;Rosa-mTmG mice received tamoxifen once daily, 0.5 mg per day for 5 consecutive days, followed by the indicated washout period before bleomycin injury and ASO treatment. GFP-labelled Sftpc-lineage cells were analysed by immunofluorescence, PCLS live imaging or cell sorting.

### Primary murine ATII cell isolation and culture

Murine Alveolar type II cells were isolated from adult lung of C57BL/6 of 8 weeks. Lungs were perfused with 10 ml of PBS from right ventricle and digested with 1ml of Dispase (SIAL-corning-354235) injected intra-tracheally and immerse in 2 ml of Dispase for 30 min at room temperature. Then, the distal part of lungs was mechanically dissociated using a McIlwain tissue chopper (Metrohm, USA) and treated with 20 µg/ml DNAse II (Sigma-Aldrich-04536282001) for 10 min at 37 °C. Cell suspension was filtered using a 100 µm-pore size cell strainer (starlab-CC8111-0102) in order to remove ATI cells and then using a 40 µm-pore size cell strainer (starlab-CC8111-0042) for promoting single cell suspension and centrifuged at 400 × g for 10 min. The pellet was then resuspended in Dulbecco’s Mem with Glutamax I (Gibco) (Dulbecco’s modified Eagle’s medium with glutamine, sodium pyruvate, pyridoxine and 4.5 g/l glucose-31966-021) supplemented with 10% FBS and antibiotics antibiotic antimycotic Penicillin and Streptomycin in order to promote negative selection of mesenchymal cells (fibroblasts) and macrophages by 2 differential adherences on plastic culture dishes of 1 h. Then, cells were centrifuged at 400 × g for 10 min and resuspended in a solution containing PBS/0,5% Bovine Serum Albumin (BSA, Sigma-A2153-100G) and 0,5 M EDTA. Following, immunity cells were removed by negative selection using simultaneous incubation with a Dynabeads Untouched Mouse T Cells Kit (Invitrogen-11413D) and Dynabeads mouse DC Enrichment Kit (Invitrogen-11429D). Recovered cells were grown on plates previously coated with Fibronectin 1mg/mL (Life technologies-33010018) and Bovin Gelatin 0.2% (Sigma Aldrich-G9391) in order to promote the ATII to ATI trans-differentiation, in Pneuma-Cult medium (Stem Cell-005001) supplemented with 10% foetal bovine serum (FBS) (Euroclone) and antibiotic antimycotic Penicillin and Streptomycin solution, according to the manufacturer’s instructions.

### Primary murine lung fibroblast isolation and culture

Primary murine lung fibroblasts were isolated from adult lung of C57BL/6 mice at 8 weeks. Lungs were perfused with 10 ml of PBS from right ventricle and mechanically dissociated with scissors. Tissues were digested incubating with an enzyme mix following the manufacturer’s instruction (Skeletal muscle dissociation kit, Miltenyi Biotec-130-098-305), for 30 min at 37 °C in agitation. The solution was filtered a 70 µm-pore size cell strainer (Falcon-CLS352350) and centrifuged at 400 × g for 10 min. The pellet was resuspended in Dulbecco’s Mem with Glutamax I (Gibco) (Dulbecco’s modified Eagle’s medium with glutamine, sodium pyruvate, pyridoxine and 4.5 g/l glucose-31966-021) supplemented with 10% FBS and antibiotics antibiotic antimycotic Penicillin and Streptomycin and incubated at 37 °C, 5% CO2 for 4 h in order to allow the attachment of fibroblasts. After 4 h medium was changed to remove macrophages and endothelial cells. Enzymes used for tissue digestion: 200 µl of Enzyme D, 50 µl of Enzyme P, 36 µl of Enzyme A in 4.70 ml of DMEM (pre-warmed at 37 °C).

### Human samples

#### Human ATII cells

Lungs were obtained from control organ donors without evidence of respiratory disease through the Gift of Life Donor Program (Philadelphia, PA) and from IPF transplants (Temple University). ATII cells were isolated by tissue digestion with elastase, followed by mechanical dissociation, separation using a density gradient, and purification by magnetic microbeads, as we previously described ^58^. This study was performed in accordance with the Declaration of Helsinki and approved by the IRB at Temple University. Informed written consent was obtained.

#### Human Lung Histological samples

Lung tissues from patients with IPF were obtained from explanted lungs obtained at the time of transplantation. All patients provided written informed consent and the study was approved by the University of Michigan Institutional Review Board, Ann Arbor, MI, USA (HUM00105694). Diagnoses of patients with IPF were established by clinical pathological criteria and confirmed by multidisciplinary consensus conference. Normal control lungs were obtained from deceased donors whose lungs were deemed unsuitable for transplant and were provided by Gift of Life, Michigan, with consent from family for tissue to be used for research purposes. No compensation was provided to subjects or family for either IPF patient samples or normal control lungs.

### miRNA mimic and ASO transfection

Primary murine and human ATII cells, primary murine lung fibroblasts and A549 cells were reverse-transfected with the indicated miRNA mimics using Lipofectamine RNAiMAX (Lipofectamine RNAiMAX, Life Technologies-1300017588) in Opti-MEM (Life Technologies-31985070). Final mimic concentrations were 30 nM for primary ATII cells, 12.5 nM for fibroblasts and 25 nM for A549 cells. cel-miR-231-3p served as a negative control and siUBC as a positive transfection control where indicated. ASO miRNA inhibitors were reverse-transfected at 30 nM into primary murine or human ATII cells, primary lung fibroblasts and A549 cells; a non-targeting ASO served as control. Cells were analysed 72 h after transfection unless otherwise specified.

### Re-analysis of a pre-existing phenotypic miRNA screen

Results of an arrayed phenotypic screening dataset, previously generated in our laboratory ^31^, comprising 2,042 human mature miRNA mimics tested in primary murine ATII cells was newly interrogated in the present study for the combined occurrence of impaired ATII-to-ATI differentiation and increased p21 positivity. RAGE-positive cells were used as a phenotypic readout of ATI differentiation, nuclear p21 as a senescence-associated readout, and total cell number as an assay-quality/toxicity readout. The screening dataset was used for candidate prioritization; selected miRNAs were subsequently evaluated in independently performed validation experiments. Details of the original screening platform are provided in Volpe, Zandomenego et al.^31^.

### Conditioned-medium transfer assays

For ATII-to-fibroblast transfer, primary murine ATII cells were transfected with the indicated miRNA mimics. After 72 h, conditioned medium was collected and filtered through molecular-weight-cutoff centrifugal filters as specified for each experiment (50-10kDa, 10-3kDa and <3kDa) (Merck life science-UFC501096, UFC500396, UFC505024). Primary lung fibroblasts were exposed for 72 h to 50 µl conditioned medium diluted in 100 µl high-glucose DMEM containing 0.5% FBS, then fixed in 4% PFA for immunofluorescence. For reciprocal fibroblast-to-ATII transfer, conditioned medium was collected 72 h after fibroblast transfection, filtered as indicated and transferred to primary ATII cells (50 µl conditioned medium in 100 µl PneumaCult medium containing 10% FBS) (Voden-05050). ATII cells were analysed after 4 days.

### PCLS preparation, live imaging and Fiji-based video analysis

Lungs for the precision cut lung slices were harvested from SftpcCreERT2/ROSAmTmG transgenic mice of 8 weeks upon the treatment with tamoxifen (0.5 mg per day for 5 consecutive days). Lungs were perfused with 10 ml of PBS from right ventricle and 1ml of Low Melting Agarose 1.5% (Life technologies-R0801) in Dulbecco’s Mem with Glutamax I (Gibco) (Dulbecco’s modified Eagle’s medium with glutamine, sodium pyruvate, pyridoxine and 4.5 g/l glucose-31966-021) is injected intra-tracheally. Then, lung was put in a petri dish with PBS in ice to allow the solidification of low melting agarose. After 30 minutes in ice, left lobe was selected and excised to be encapsulated into a cylinder of agarose 3% in PBS. Vibratome (Alabama R&D Tissue Slicer) was used to perform lung sections of 150 µm of thickness, using a frequency of 100 Hz and a speed of blade of 3-12 µm/s. Slices were transferred into 12-well plate filled with cultivation medium (Dulbecco’s modified Eagle’s medium with glutamine, sodium pyruvate, pyridoxine and 4.5 g/l glucose-31966-021) supplemented with 10% foetal bovine serum (FBS) (Euroclone) and antibiotic antimycotic Penicillin and Streptomycin solution, according to the manufacturer’s instructions. Medium was changed every hour on the first day to dissolve and eliminate the excess agarose. The next day, precision-cut lung slices were stained and acquired using the Operetta CLS (Revvity). For the regenerative-dynamics experiment, slices were maintained in culture for 6 days and live-imaged for 12 h beginning 24 h after isolation, with acquisitions every 3 h using the Operetta CLS high-content imaging system (Revvity). Time-lapse sequences were analysed in Fiji/ImageJ using TrackMate plugin. Image sequences were calibrated using acquisition metadata and evaluated in the GFP channel to follow lineage-labelled Sftpc-derived epithelial cells over time. Individual GFP-positive cells that remained identifiable across the imaging sequence were followed between successive time points to quantify cell displacement. Net displacement was calculated from the change in cell-centroid position between the first and last analysed frames. Morphological maturation was assessed from GFP-positive cell shape by outlining individual cells and quantifying elongation from major-and minor-axis measurements; the same analysis settings and classification criterion were applied to all experimental groups. Endpoint GFP-positive cell abundance and collagen-associated immunofluorescence were quantified from matched fields using identical acquisition and analysis settings across conditions.

### Immunofluorescence, histology and miRNA in situ hybridization

Cultured cells, frozen lung sections and FFPE lung sections were processed for immunofluorescence using standard fixation, permeabilization, blocking and antibody-incubation procedures. Nuclei were counterstained with Hoechst 33342. FFPE sections underwent heat-mediated antigen retrieval before staining. Fluorescent in situ hybridization for mature miRNAs was performed on FFPE lung sections with biotin-conjugated LNA probes (Qiagen-339111), streptavidin-HRP (merk like science-GERPN1231) detection and TSA-Cy5 amplification (Akoya-Aurogene-NEL745001KT, followed by immunofluorescence where indicated. Masson’s trichrome and haematoxylin and eosin staining were performed on paraffin sections using commercial reagents. Antibodies and probe sequences are listed in Tables 1-3.

**Table 1:**
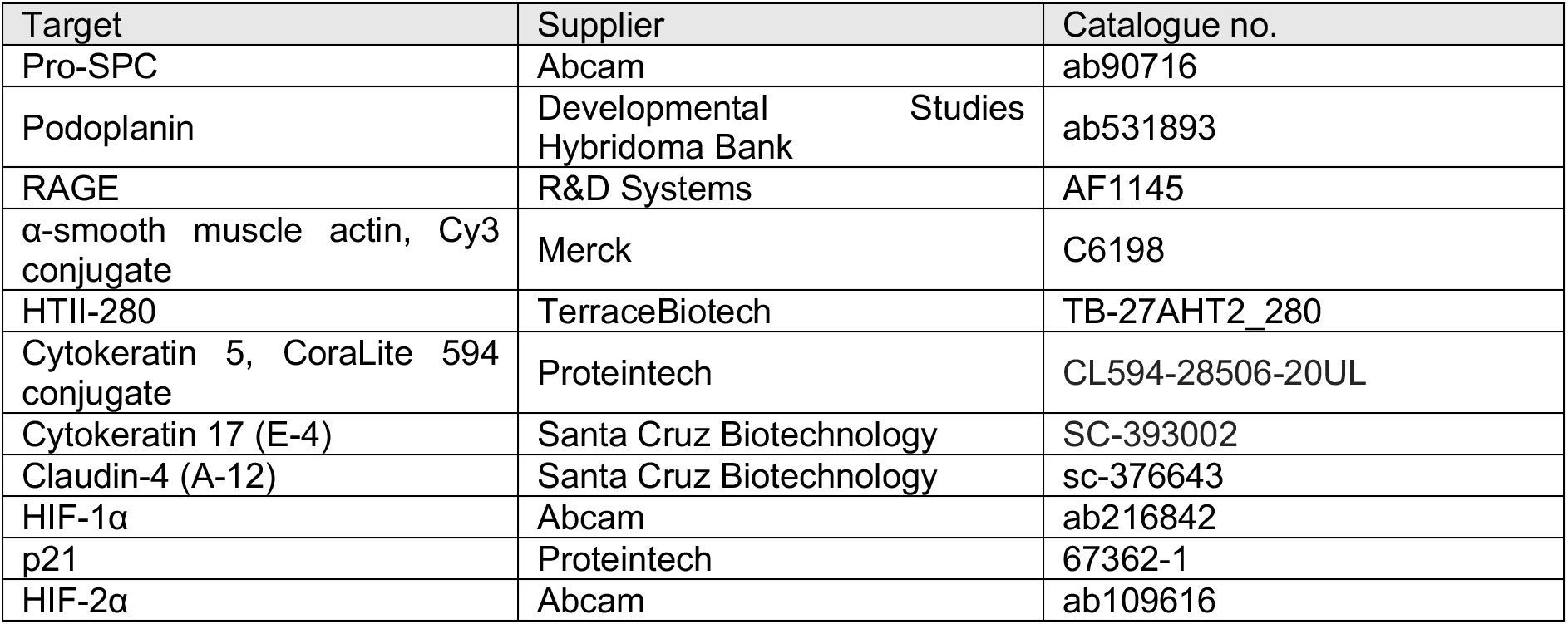
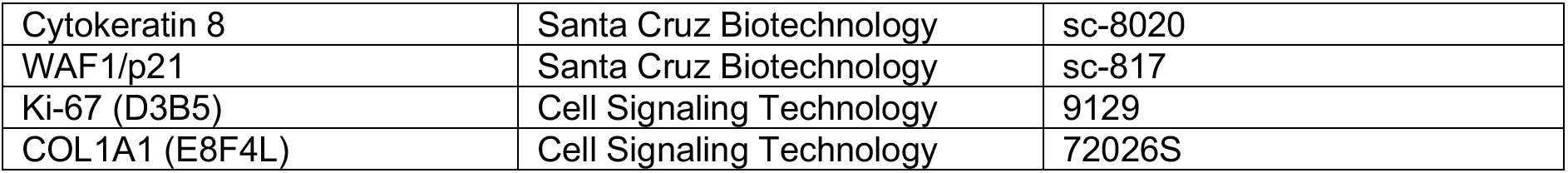
Primary antibodies.

**Table 2:** Secondary antibodies.

| Reagent | Supplier | Catalogue no. |
| --- | --- | --- |
| Alexa Fluor 568 donkey anti-goat IgG | Invitrogen | A-11057 |
| Alexa Fluor 488 donkey anti-goat IgG | Invitrogen | A-11057 |
| Alexa Fluor 647 donkey anti-goat IgG | Invitrogen | A-21447 |
| Alexa Fluor 568 donkey anti-mouse IgG | Invitrogen | A-10037 |
| Alexa Fluor 488 donkey anti-mouse IgG | Invitrogen | A-21202 |
| Alexa Fluor 647 donkey anti-mouse IgG | Invitrogen | A-31571 |
| Alexa Fluor 594 donkey anti-rabbit IgG | Invitrogen | A-21207 |
| Alexa Fluor 488 donkey anti-rabbit IgG | Invitrogen | A-21206 |
| Alexa Fluor 647 donkey anti-rabbit IgG | Invitrogen | A-31576 |
| Alexa Fluor 488 goat anti-Syrian hamster IgG H&L | Abcam | ab180063 |
| Alexa Fluor 647 goat anti-Syrian hamster IgG H&L | Abcam | ab180117 |

**Table 3:**
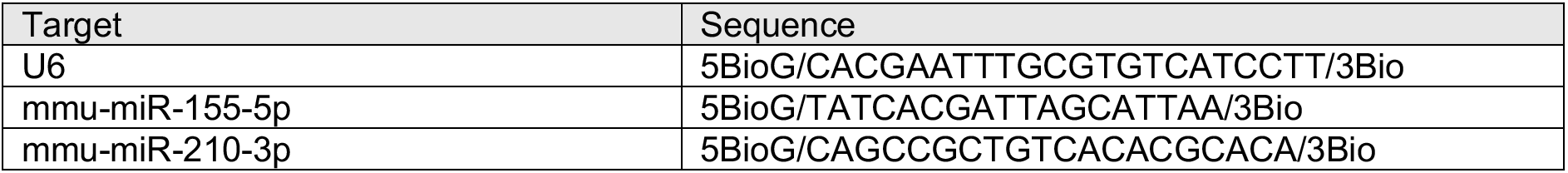
LNA probe sequences.

### RNA isolation and quantitative PCR

Total RNA was isolated using miRNeasy Mini or Micro kits (Qiagen-217084), depending on sample type. For mRNA analysis, 500 ng total RNA was reverse-transcribed using M-MLV reverse transcriptase (Life technologies-28025-013) and analysed by SYBR Green (Life technologies-S33102 quantitative PCR on a CFX96 Real-Time System (Bio-Rad). Mature miRNAs were reverse-transcribed using the miRCURY LNA RT Kit (Qiagen-339340) and quantified using miRCURY LNA miRNA PCR assays (Qiagen-339346) and miRCURY SYBR Green Master Mix. Primer sequences are provided in Table 4.

**Table 4:**
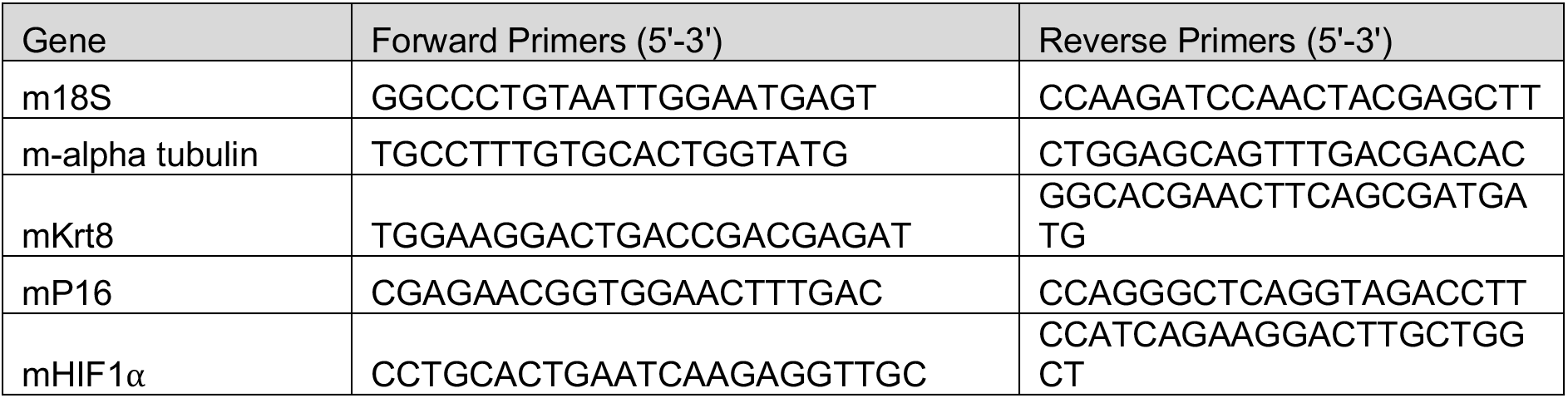
Sequences of Primers.

### Hydroxyproline assay

Lung hydroxyproline was quantified using the Hydroxyproline Assay Kit (Vinci Biochem-MA-0101). Wet lung tissue (10 mg) was homogenized in 100 µl water and hydrolysed with an equal volume of 12 M HCl for 3 h at 120 °C. Supernatants were evaporated at 60 °C and processed with chloramine T and DMAB reagents according to the manufacturer’s instructions. Absorbance was measured at 560 nm and hydroxyproline content was calculated from a standard curve.

### Flow cytometry

Single-cell lung suspensions were generated by Dispase-based dissociation and differential adherence. Cells were stained with Fc block, a fixable viability dye and antibodies against CD45, CD31, EpCAM, MHCII and SPC. Uptake of fluorescently labelled ASO was quantified within the indicated populations. Samples were acquired on a BD FACSAria II. Unstained, single-colour compensation and isotype controls were included. Flow-cytometry antibodies are listed in Table 5.

**Table 5:** Flow-cytometry antibodies.

| Marker/fluorophore | Supplier | Clone/isotype or catalogue |
| --- | --- | --- |
| CD45-APC-Cy7 | BD Pharmingen | IgG2b, κ, 561037 |
| CD31-BV421 | BD Pharmingen | IgG2a, κ, 563356 |
| EpCAM-BV650 | BioLegend | IgG2b, κ, 118241 |
| MHCII-BV711 | BioLegend | IgG2b, κ, 107643 |
| MHCII-Alexa700 | BioLegend | IgG2b, κ, 107622 |

### RNA sequencing and hierarchical clustering analysis

Lineage-traced ATII cells were isolated from bleomycin-treated Sftpc-CreERT2;Rosa-mTmG mice following therapeutic administration of ASO-miR-155-5p or ASO-miR-210-3p. Total RNA was subjected to RNA sequencing, and gene expression was quantified for 45,777 annotated mouse transcripts, generating raw read counts, FPKM and TPM values across six samples (BLEO2, BLEO3, ASO155-2, ASO155-3, ASO210-1 and ASO210-2). Differential expression analysis was performed using DESeq2 with a three-group design (bleomycin, ASO-miR-155-5p and ASO-miR-210-3p). Genes with undetectable or near-zero expression across samples were removed before differential expression analysis, for the selected genes statistics were computed for two contrasts: ASO-miR-155-5p versus bleomycin and ASO-miR-210-3p.

For two-way hierarchical clustering, the gene set was defined as the union of protein-coding genes meeting the differential-expression criteria (|fold change| > 1 and raw P < 0.05) in either ASO-miR-155-5p versus bleomycin or ASO-miR-210-3p versus bleomycin comparison. Genes differentially expressed exclusively between the two ASO treatments were excluded. DESeq2 size-factor-normalized counts were extracted for the six samples. For genes represented by multiple transcript entries, the entry with the highest mean normalized expression across samples was retained. Normalized counts were transformed as log2(count + 1) and standardized gene-wise as Z-scores across the six samples. Two-way hierarchical clustering of genes and samples was performed using Euclidean distance and complete linkage, and the resulting relative expression patterns were visualized as a clustered heatmap.

### Pathway analysis

Of the 21,352 genes for which differential expression statistics were computed, genes were selected separately for the ASO-miR-155-5p versus bleomycin and ASO-miR-210-3p versus bleomycin contrasts using the following criteria: protein-coding biotype, fold change ≥1 or ≤−1, and raw P ≤ 0.05. The resulting lists were independently submitted to Ingenuity Pathway Analysis (IPA). Canonical pathway enrichment was performed in IPA using a right-tailed Fisher’s exact test. Pathways with P < 0.05 were considered enriched, and predicted activation state was inferred from the IPA activation z-score. Canonical pathways identified by IPA were subsequently filtered to enrich for programmes jointly regulated by both ASOs and containing conserved candidate miRNA targets. Pathway molecules were intersected with conserved derepressed miR-155-5p and miR-210-3p targets, retaining pathways containing ≥2 targets (n = 35). These were further restricted to pathways containing ≥5 genes significantly regulated in both ASOs-versus-bleomycin comparisons, yielding 24 pathways for downstream analysis. Robustness to miRNA target definition was assessed by recalculating target enrichment across the 24 pathways using human-only, mouse-only or conserved human–mouse target predictions, in each case restricted to targets derepressed following ASO treatment. Pathway rankings were compared using pairwise Spearman correlations and Kendall’s coefficient of concordance, showing consistent rankings across target definitions (Spearman ρ = 0.624–0.695, all P < 0.001; Kendall’s W = 0.553, P = 0.024). Pathways were classified according to the directionality of their shared differentially expressed genes using a two-sided exact binomial test against an equal proportion of up- and downregulated genes (P_up = 0.5). Pathways with P < 0.05 and >50% upregulated genes were classified as direct-target driven, those with P < 0.05 and <50% upregulated genes as downstream effectors, and those without significant directional bias (P ≥ 0.05) as bidirectionally remodelled. This identified 4 direct-target-driven, 5 downstream-effector and 15 bidirectionally remodelled pathways.

### Public single-cell RNA-sequencing analysis

Public lung scRNA-seq datasets were interrogated in BBrowserX (BioTuring Inc.) to assess conservation of the ASO-restored transcriptional programme. Differential expression between fibrotic and control alveolar epithelial populations was calculated using the built-in pseudobulk DESeq2 workflow, requiring expression coverage >5% and FDR <0.05. The analysis included human datasets from Habermann et al. (GSE135893), Adams et al. (GSE136831) and Yao et al. (GSE146981), and murine datasets from Peyser et al. (GSE129605), Parimon et al. (GSE134741) and Strunz et al. (GSE141259), as used in the final cross-dataset analysis. Genes were aligned by gene symbol and compared according to the direction of differential expression in fibrosis versus ASO treatment. The Habermann dataset was further analysed for 30 selected miRNAs predicted target in epithelial/mesenchymal-compartment while retaining the original author-provided annotations and UMAP coordinates.

### Statistical analysis

Statistical analyses were performed in GraphPad Prism. Data are presented as mean ± s.d. unless otherwise indicated. Individual points represent independent biological replicates for in vitro experiments or individual animals for in vivo experiments; where a plotted point is the mean of technical replicate wells, this is stated in the corresponding figure legend. Comparisons among multiple groups were performed using one- or two-way ANOVA followed by the prespecified multiple-comparisons test. Dunnett’s test was used for comparisons of multiple groups with a single control, Tukey’s test for all pairwise comparisons where indicated, and Šídák’s test for selected pairwise comparisons. Two-group comparisons were performed using an unpaired two-tailed Welch’s t-test unless otherwise stated. P < 0.05 was considered statistically significant; exact P values are reported where appropriate.

## Data availability

Processed bulk RNA-sequencing data generated in this study will be made publicly available in an appropriate repository upon publication. Public scRNA-seq datasets used in this study are available under GSE135893, GSE136831, GSE146981, GSE129605, GSE134741 and GSE141259. Source Data will accompany the manuscript.

## Acknowledgments

We are grateful to Prof. Serena Zacchigna for generously providing the ROSAmT/mG mice. We also thank Dr. Simone Vodret for assistance with animal procedures, and Prof. Giovanni Sorrentino and Dr. Beatrice Anfuso for providing access to instrumentation for precision-cut lung slices (PCLS) generation. This work was supported by the Fondazione CRTrieste, the Beneficentia Stiftung Foundation, and Associazione AMAR-FVG.

## Authors contribution

L.B. conceived the study, with contributions from G.Z. and M.C. G.Z. and M.T. performed primary cell isolations, in vitro cellular experiments and in vivo experiments. L.B., G.Z., A.M.D.I and D.L. performed gene expression and bioinformatic analyses. B.K. and K.B. provided primary human ATII cells Human from lungs were obtained through the Lung Biobank at Temple University. A.A.R. performed experiments on human histological samples under the supervision of G.L. L.B. and G.Z. analysed and interpreted the data. L.B. and G.Z. wrote the original draft of the manuscript. P.C., F.S. and M.C.V. contributed to data interpretation and provided critical intellectual input. M.C. and G.L. provided scientific guidance and supervision. L.B. provided overall supervision and directed the study. All authors contributed to the critical revision of the manuscript and approved the final version.

## Declaration of generative AI and AI-assisted technologies in the manuscript preparation process

During the preparation of this work, the authors used ChatGPT Edu (OpenAI) to assist with English-language editing and to improve the clarity and readability of the manuscript. Claude for Science (Anthropic) was used to assist with RNA-sequencing data analysis, the classification and annotation of IPA-derived pathways, and their integration with predicted miRNA targets. Claude (Anthropic) and ChatGPT Edu (OpenAI) were also used to assist with the preparation of the graphical abstract. All AI-assisted outputs were critically reviewed, verified and, where appropriate, edited by the authors. The authors take full responsibility for the analyses, interpretation, conclusions and content of the manuscript.

## Extended Data Figure

**Extended Data Fig. 1:**
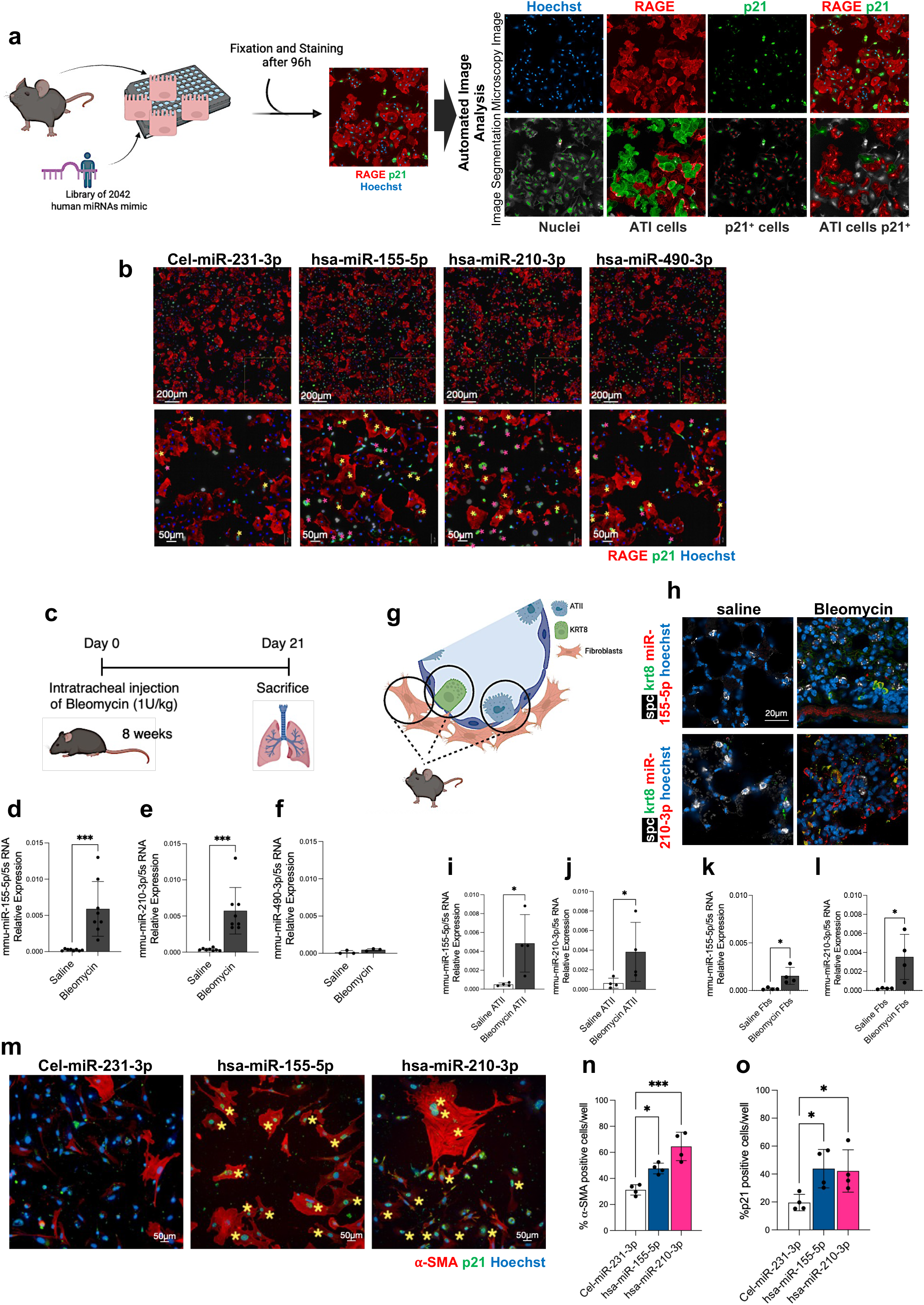
Phenotypic validation of miR-155-5p and miR-210-3p as regulators of epithelial differentiation, senescence and fibroblast activation. **a**) Schematic representation of the high-content phenotypic screening workflow. Primary murine ATII cells were transfected with a library of 2,042 human miRNA mimics, fixed after 96 h and stained for RAGE, p21 and Hoechst. Representative images illustrate the automated image-analysis pipeline used to segment nuclei and identify RAGE⁺ ATI cells, p21⁺ cells and RAGE⁺p21⁺ cells. **b**) Representative immunofluorescence images of primary ATII cells transfected with cel-miR-231-3p, hsa-miR-155-5p, hsa-miR-210-3p or hsa-miR-490-3p and stained for RAGE, p21 and Hoechst. Asterisks indicate representative p21-positive cells. (pink=Fibroblasts and yellow=ATII cells) **c**) Experimental design of the bleomycin-induced pulmonary fibrosis model used for analysis at day 21. **d–f**) Quantification of miRNA expression levels in saline- and bleomycin-treated lungs. **g**) Schematic representation of the alveolar niche analysed in bleomycin-treated lungs, including ATII, ATI, KRT8⁺ transitional epithelial cells and fibroblasts. **h**) Representative combined miRNA in situ hybridization and immunofluorescence images showing miR-155-5p or miR-210-3p (red) together with SPC (white) and KRT8 (green) in saline- and bleomycin-treated lungs. **i–j**) Quantification of miR-155-5p- and miR-210-3p-expression levels in murine ATII cells healthy vs bleomycin treated mice. **k–l**) Quantification of miR-155-5p- and miR-210-3p-expression levels in murine fibroblasts healthy vs bleomycin treated mice. m) Representative immunofluorescence images of primary lung fibroblasts transfected with cel-miR-231-3p, hsa-miR-155-5p or hsa-miR-210-3p and stained for α-SMA (red), p21(green) and Hoechst(blue). Asterisks indicate representative p21-positive cells. **n, o**) Quantification of α-SMA-positive (**n**) and p21-positive (**o**) fibroblasts following miRNA transfection. Statistical significance was determined using unpaired t test for d, e, f, i, j, k, l and One-way ANOVA followed by Dunnett’s for n, o with multiple-comparison correction where appropriate. *P < 0.05; ***P < 0.001.

**Extended Data Fig. 2:**
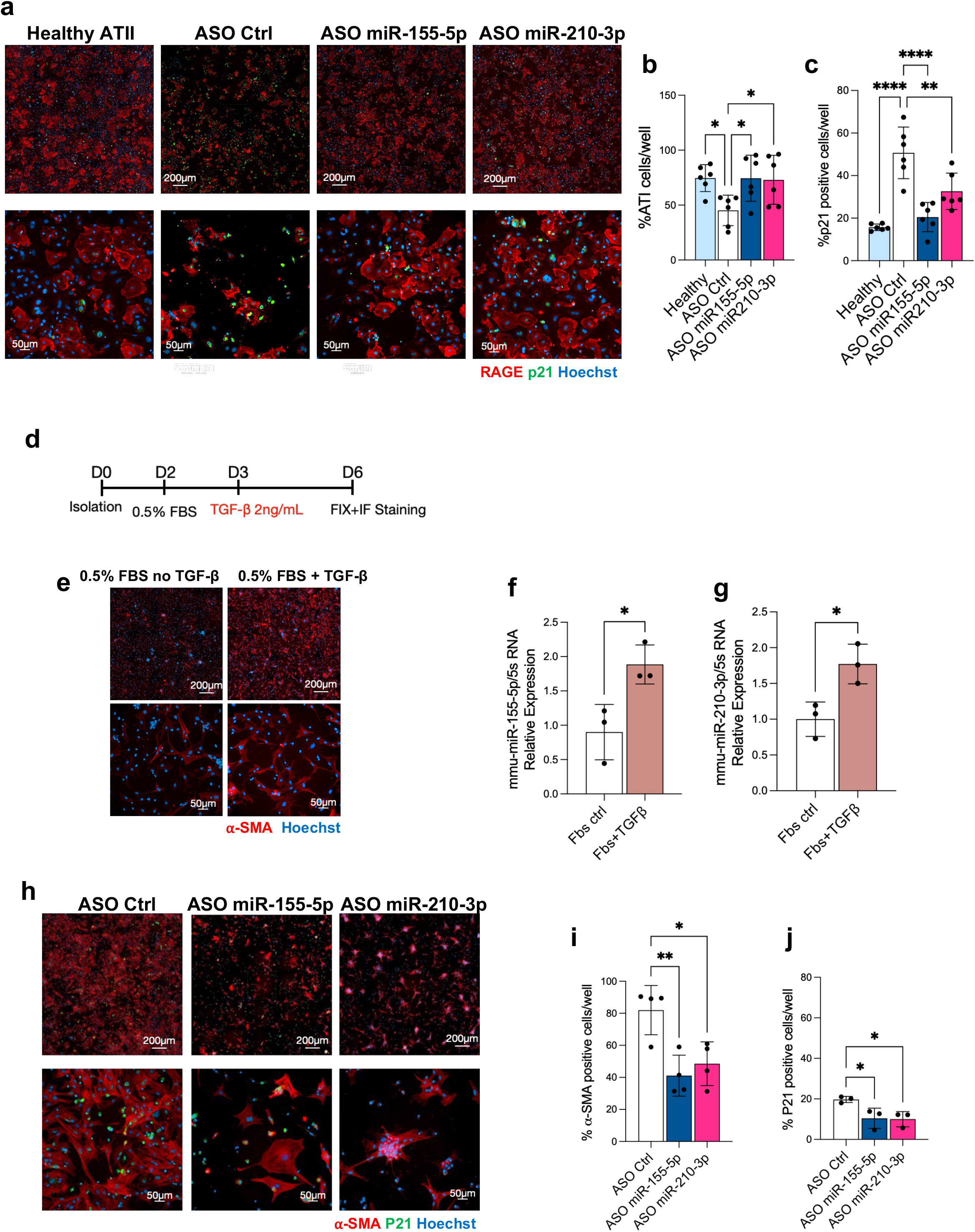
ASO-mediated inhibition of miR-155-5p and miR-210-3p restores ATII differentiation and attenuates fibroblast activation. **a)** Representative immunofluorescence images of primary murine ATII cells isolated from healthy mice and after 21 days from bleomycin injection. Bleomycin ATII cells were treated with control ASO, ASO-miR-155-5p or ASO-miR-210-3p as well as healthy ATII cells and stained for RAGE (red), p21 (green) and Hoechst (blue). **b, c**) Quantification of RAGE-positive ATI cells (**b**) and p21-positive cells (**c**) following ASO treatment. **d**) Experimental design of TGF-β-induced activation of primary lung fibroblasts. Cells were maintained in 0.5% FBS and stimulated with TGF-β (2 ng ml⁻¹) before fixation and immunofluorescence analysis. **e**) Representative α-SMA (red) immunofluorescence images of fibroblasts maintained in 0.5% FBS in the absence or presence of TGF-β. **f, g**) Quantification of miRNAs expression levels after fibroblast activation following TGF-β stimulation. **h**) Representative immunofluorescence images of activated fibroblasts treated with control ASO, ASO-miR-155-5p or ASO-miR-210-3p and stained for α-SMA (red), p21 (green) and Hoechst (blue). **i, j**) Quantification of α-SMA-positive (**i**) and p21-positive (**j**) fibroblasts following hypoxamiR inhibition. Scale bars are indicated in the images. Statistical significance was determined using unpaired t test for f, g and one-way ANOVA followed by Dunnett’s multiple-comparisons test for b, c, I, j with multiple-comparison correction where appropriate. *P < 0.05; **P < 0.01; ***P < 0.001.

**Extended Data Fig. 3:**
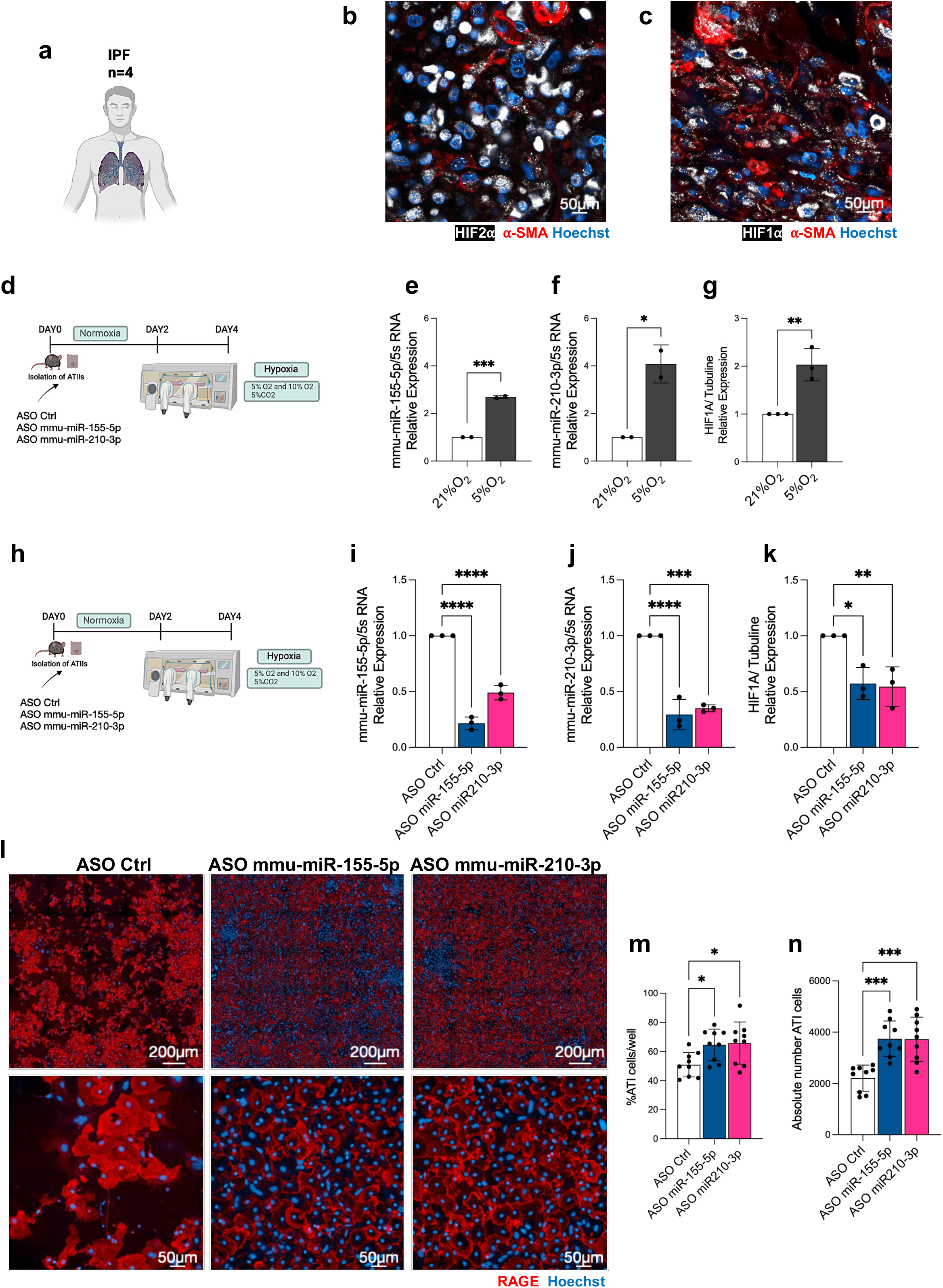
Hypoxia induces miR-155-5p and miR-210-3p and ASO-mediated inhibition disrupts the HIF–hypoxamiR axis. **a)** Schematic representation of the human IPF lung samples analyzed (n = 4). **b, c)** Representative immunofluorescence images of human IPF lungs stained for (**b**) HIF-2α (white) and α-SMA(red) or (**c**) HIF-1α (white) and α-SMA (red), with Hoechst (blue) nuclear counterstaining. **d)** Experimental design of primary ATII cell exposure to hypoxia. Cells were maintained under normoxia before exposure to reduced oxygen tension for 48 h. **e, f)** Relative expression of miR-155-5p and miR-210-3p in primary ATII cells cultured under normoxic or hypoxic conditions. **g)** Quantification of HIF1A expression levels following hypoxic exposure. **h)** Experimental design of ASO treatment under hypoxic conditions. Primary ATII cells were treated with control ASO, ASO-miR-155-5p or ASO-miR-210-3p before exposure to hypoxia. **i, j)** Quantification of miRNAs expression levels, miR-155-5p (**i**) and miR-210-3p (**j**). **k)** Quantification of HIF1A expression levels following ASO treatment under hypoxic conditions. **l)** Representative immunofluorescence images of hypoxic ATII cells treated with control ASO, ASO-miR-155-5p or ASO-miR-210-3p stained for RAGE (red) and Hoechst (blue) nuclear counterstaining **m,n)** Quantification of RAGE-positive ATI differentiation (**m**) and total cell number/survival (**n**) following ASO treatment. Nuclei were counterstained with Hoechst. Statistical significance was determined using unpaired t test for e,f,g and One-way ANOVA followed by Dunnett’s for i,j,k,m,n with multiple-comparison correction where appropriate. *P < 0.05; **P < 0.01; ***P < 0.001; ****P < 0.0001.

**Extended Data Fig. 4:**
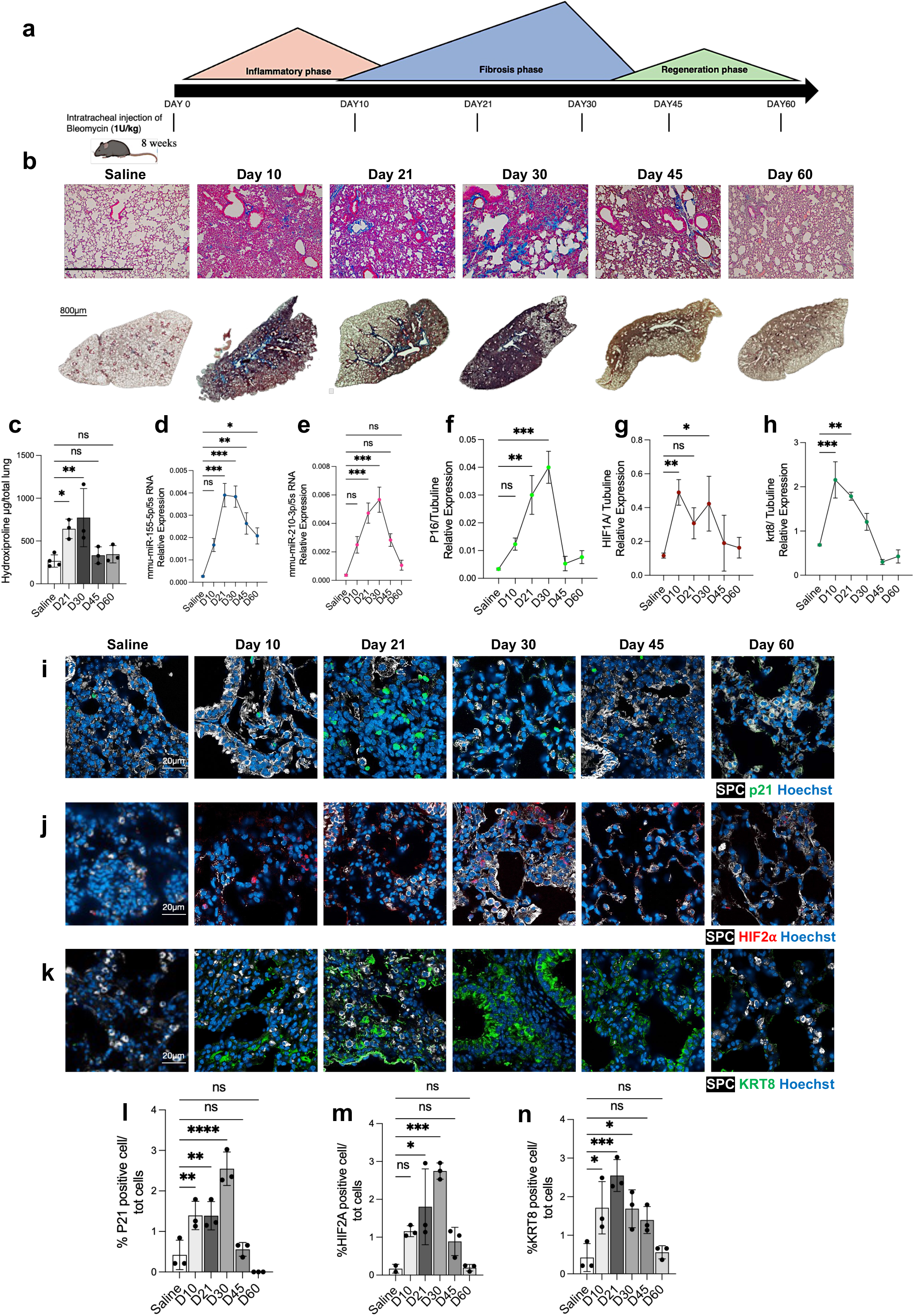
miR-155-5p and miR-210-3p dynamically track fibrosis progression and resolution following bleomycin injury. **a)** Schematic representation of the temporal evolution of bleomycin-induced lung injury, encompassing inflammatory, fibrotic and regenerative phases, and of the collection time points at days 10, 21, 30, 45 and 60. **b)** Representative higher-magnification and whole-lung Masson’s trichrome images from saline-treated mice and mice collected at days 10, 21, 30, 45 and 60 after bleomycin administration. **c)** hydroxyproline content across the fibrosis–resolution time course. **d, e)** miRNAs expression levels at different time points **f–h),** Relative mRNA expression of *Cdkn2a* (p16) (**f**), *Hif1a* (**g**) and *Krt8* (**h**), normalized to *Tubulin*, during fibrosis progression and resolution. **i–k)** Representative immunofluorescence images of SPC (white) together with p21 (green) (**i**), HIF-2α (red) (**j**) and KRT8 (green) (**k**) at the indicated time points. **l–n)** Quantification of p21-positive (**l**), HIF-2α-positive (**m**) and KRT8-positive (**n**) epithelial cells across the time course. Nuclei were counterstained with Hoechst (blue). Statistical significance was determined One-way ANOVA followed by Dunnett’s with multiple-comparison correction where appropriate. *P < 0.05; **P < 0.01; ***P < 0.001; ****P < 0.0001; ns, not significant.

**Extended Data Fig. 5:**
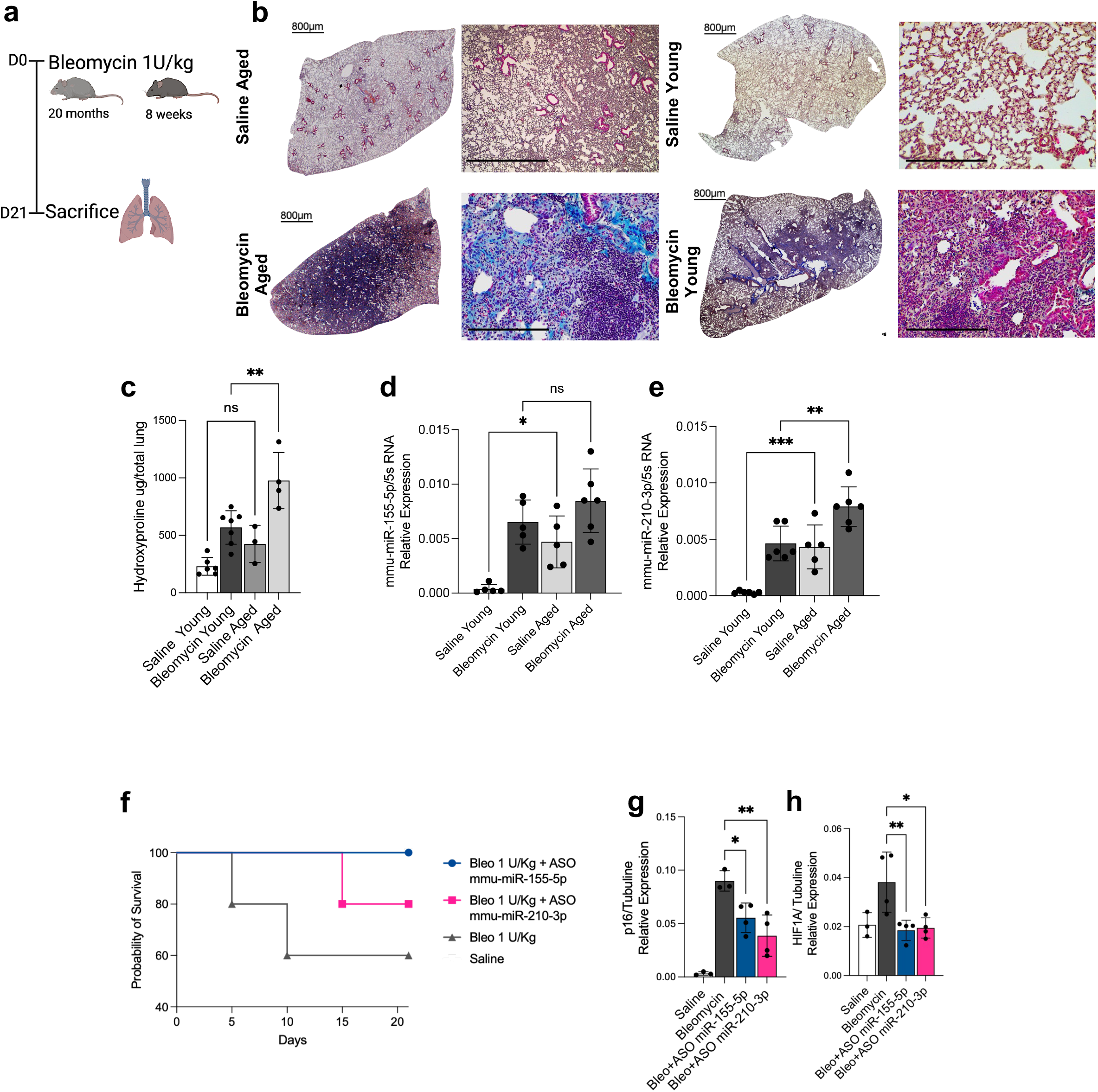
Aging exacerbates bleomycin-induced pulmonary fibrosis and impairs alveolar epithelial homeostasis. Extended Data Fig. 5 | Ageing exacerbates bleomycin-induced pulmonary fibrosis and impairs alveolar epithelial homeostasis. **a)** Experimental design comparing 20-month-old and 8-week-old mice analysed 21 days after bleomycin administration. **b)** Representative whole-lung and higher-magnification Masson’s trichrome images from saline- and bleomycin-treated aged and young mice. **c)** Lung hydroxyproline content in saline- and bleomycin-treated young and aged mice. **d,e)** Relative expression of miR-155-5p (**d**) and miR-210-3p (**e**) across age and treatment groups. **f)** Kaplan–Meier survival curves following bleomycin administration in mice treated with ASO-miR-155-5p or ASO-miR-210-3p and in saline-treated controls. **g,h)** Relative mRNA expression of p16; **g**) and *Hif1a* (**h**), normalized to tubulin, following bleomycin administration and hypoxamiR inhibition. Statistical significance was determined by one-way ANOVA followed by Dunnett’s multiple-comparisons test for **c–e,g,h**. *P < 0.05; **P < 0.01; ***P < 0.001; ns, not significant.

**Extended Data Fig. 6:**
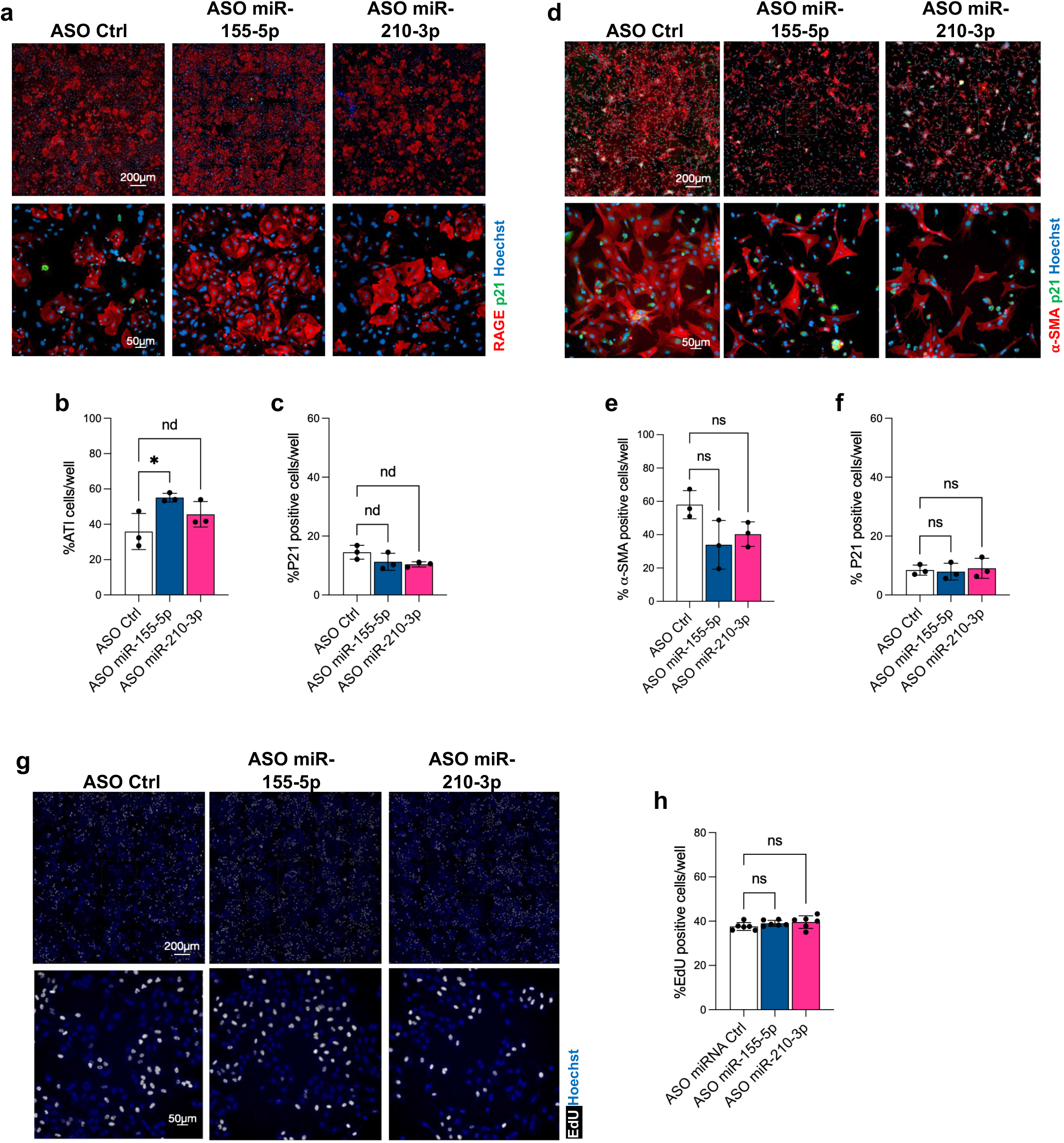
HypoxamiR inhibition does not perturb epithelial or fibroblast homeostasis or basal cell proliferation. **a)** Representative immunofluorescence images of healthy primary ATII cells treated with control ASO, ASO-miR-155-5p or ASO-miR-210-3p and stained for RAGE (red), p21 (green) and Hoechst (blue). **b, c)** Quantification of RAGE-positive (**b**) ATI differentiation and p21-positive (**c**) cells following ASO treatment. **d)** Representative immunofluorescence images of healthy primary lung fibroblasts treated with control ASO, ASO-miR-155-5p or ASO-miR-210-3p and stained for α-SMA (red), p21(green) and Hoechst (blue). **e, f)** Quantification of α-SMA-positive (**e**) and p21-positive (**f**) fibroblasts. **g)** Representative EdU (white) incorporation images from A549 cells treated with control ASO, ASO-miR-155-5p or ASO-miR-210-3p. **h)** Quantification of EdU-positive cells following ASO treatment. Statistical significance was determined One-way ANOVA followed by Dunnett’s with multiple-comparison correction where appropriate. *P < 0.05; ns, not significant.

**Extended Data Fig. 7.**
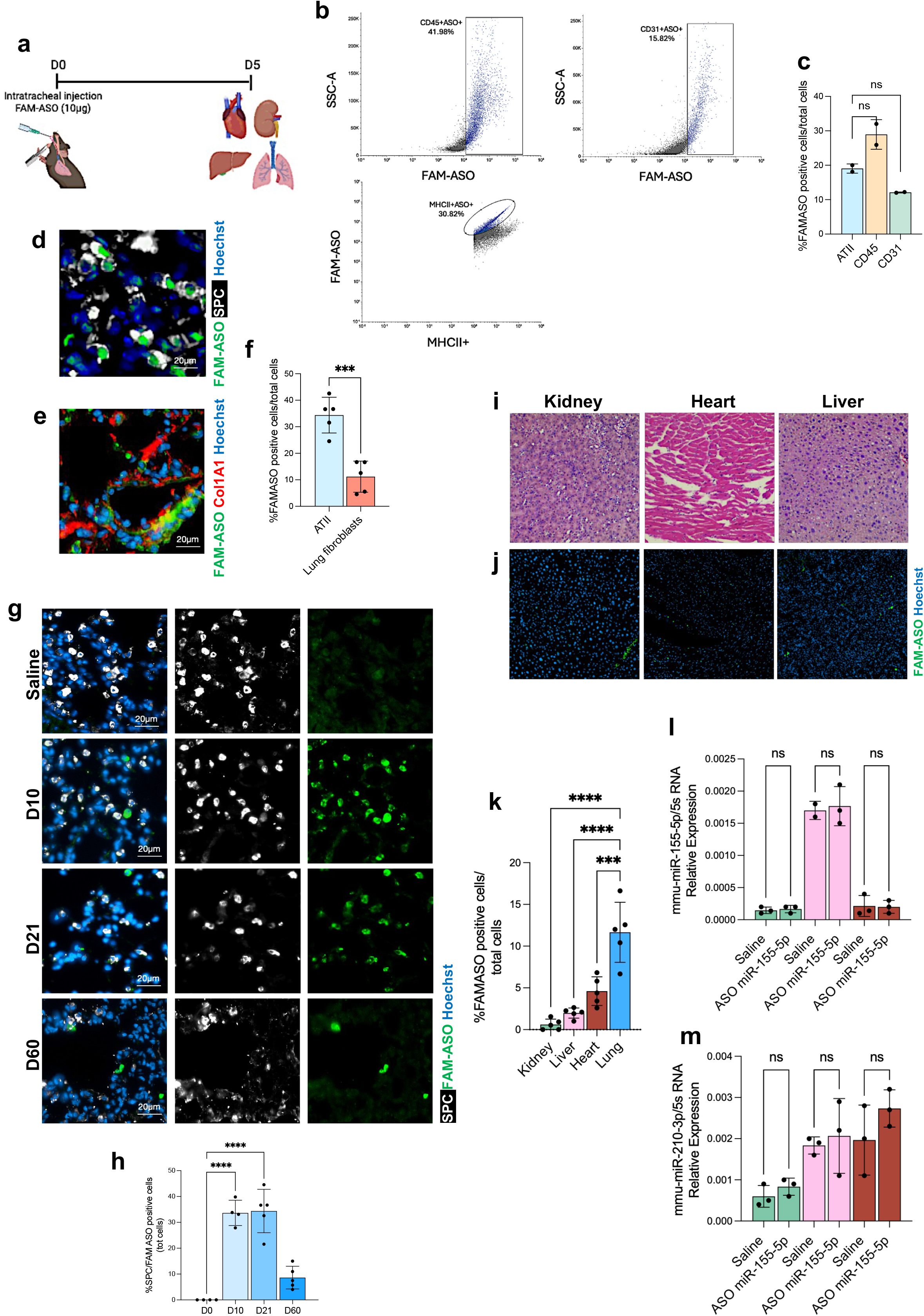
Intratracheally delivered ASOs efficiently reach the lung epithelium, persist in pulmonary tissue and show limited extrapulmonary distribution. **a)** Experimental design of FAM-labelled ASO administration and biodistribution analysis 5 days after intratracheal delivery. **b)** Representative flow-cytometric analysis of FAM-ASO uptake in the indicated pulmonary cell populations, including CD45-positive, CD31-positive and MHCII-positive cells. **c)** Quantification of FAM-ASO-positive cells across the indicated lung cell populations. **d,e)** Representative lung immunofluorescence images showing FAM-ASO (green) localization in SPC-positive alveolar epithelial cells (white; (**d**) and COL1A1-positive stromal/fibroblast cells (red) (**e**). Nuclei were counterstained with Hoechst (blue). **f)** Quantification of FAM-ASO-positive cells in ATII cells and lung fibroblasts. **g)** Representative immunofluorescence images showing pulmonary persistence of FAM-ASO (green) in SPC-positive cells (white) at days 10, 21 and 60 after administration, with saline-treated lungs as controls. Nuclei were counterstained with Hoechst (blue). **h)** Quantification of FAM-ASO-positive SPC-positive cells over time. **i,j)** Representative haematoxylin and eosin staining (**i**) and FAM-ASO fluorescence (**j**) in kidney, heart and liver following intratracheal FAM-ASO administration. Nuclei were counterstained with Hoechst (blue). **k)** Quantification of FAM-ASO-positive cells in kidney, liver, heart and lung. **l,m)** Relative expression of miR-155-5p (**l**) and miR-210-3p (**m**) in kidney, liver and heart following ASO treatment. Statistical significance was determined using unpaired t test for f and One-way ANOVA followed by Šídák’s with multiple comparisons test correction where appropriate. ***P < 0.001; ****P < 0.0001; ns, not significant

**Extended Data Fig. 8.**
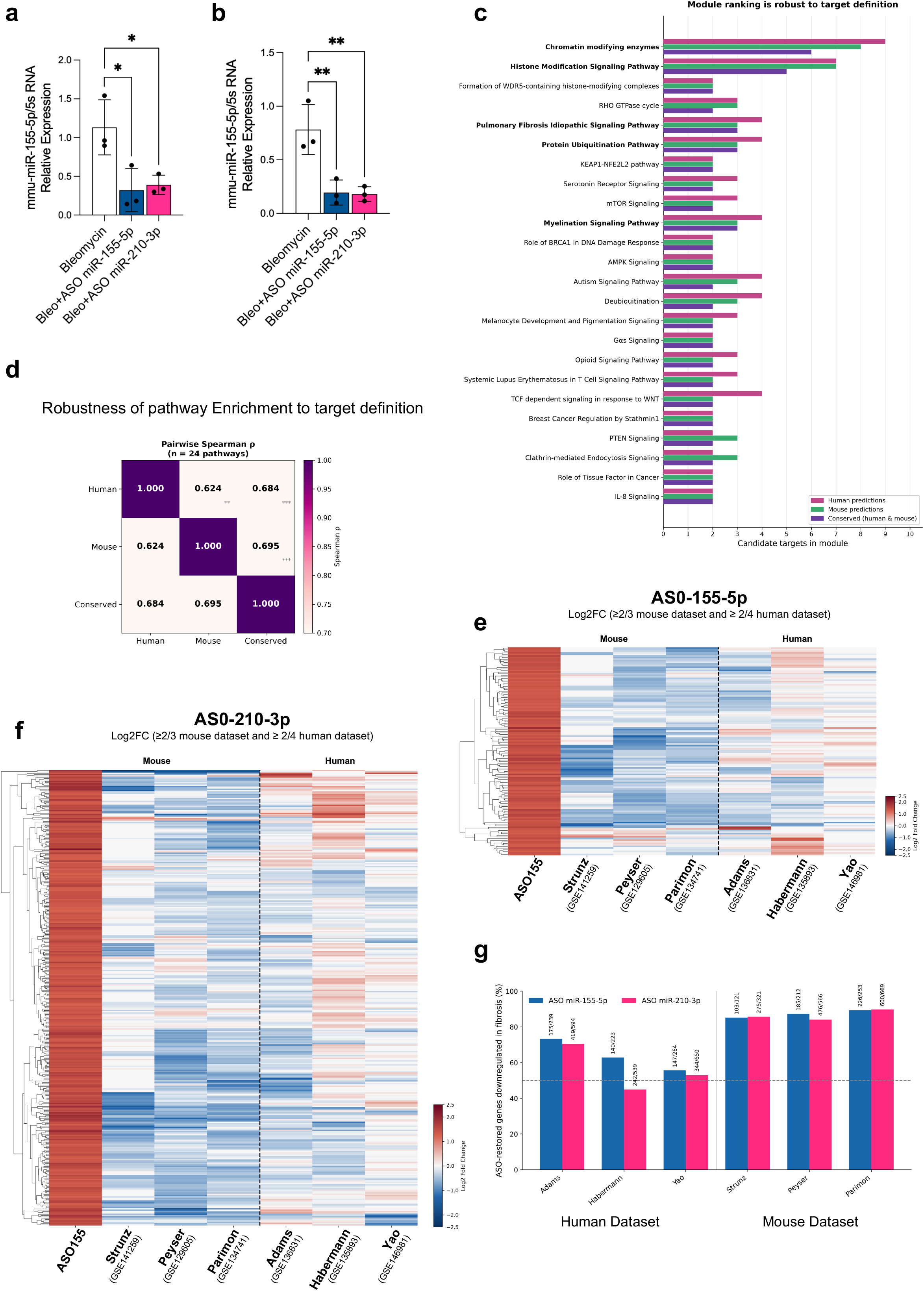
HypoxamiR inhibition restores a conserved epithelial transcriptional programme disrupted across human IPF and experimental pulmonary fibrosis. **a,b)** Quantification of miR-155-5p (**a**) and miR-210-3p (**b**) in lineage-traced ATII cells isolated from bleomycin-treated mice following therapeutic administration of ASO-miR-155-5p or ASO-miR-210-3p, confirming inhibition of the corresponding miRNA. **c)** Sensitivity analysis of pathway-module ranking using alternative definitions of candidate miRNA targets. Bars indicate the number of candidate targets assigned to each pathway module using human target predictions, mouse target predictions or targets conserved between human and mouse, demonstrating the robustness of the highest-ranking modules to target-definition strategy. **d,e)** Cross-dataset analysis of genes upregulated following ASO-miR-155-5p (**d**) or ASO-miR-210-3p (**e**) treatment in lineage-traced ATII cells and their corresponding expression changes across independent pulmonary fibrosis transcriptomic datasets. Genes upregulated following the respective ASO treatment were matched across the indicated human and murine ATII datasets; genes absent from an individual dataset are shown in white. Colours indicate the direction and magnitude of differential expression, with yellow representing increased and blue representing decreased expression. Genes were hierarchically clustered using Euclidean distance and complete linkage; dataset order was fixed. Human datasets include Adams *et al.*, Habermann *et al.* and Yao *et al.*; murine datasets include Strunz *et al.*, Peyser *et al.* and Parimon *et al. **f**)* Concordance of pathway rankings obtained using human-only, mouse-only and conserved miRNA target predictions. Values indicate Spearman’s ρ (all *P* < 0.001); overall concordance: Kendall’s W = 0.553, *P* = 0.024. **g)** Quantitative cross-dataset concordance of the transcriptional programmes restored by ASO-miR-155-5p and ASO-miR-210-3p. Bars indicate the percentage of ASO-restored genes that were reciprocally downregulated in fibrotic ATII cells across independent human (Adams, Habermann and Yao *et al.*) and murine (Strunz, Peyser and Parimon *et al.*) transcriptomic datasets. Percentages were calculated relative to the number of ASO-restored genes detected in each dataset. Values above bars indicate the number of reciprocally downregulated genes relative to the total number of detected ASO-restored genes. The dashed line indicates 50% concordance. Statistical significance in (a) was assessed by one-way ANOVA followed by Dunnett’s multiple-comparisons test. **P < 0.01; ***P < 0.001; ****P < 0.0001

**Supplementary Table 1.**
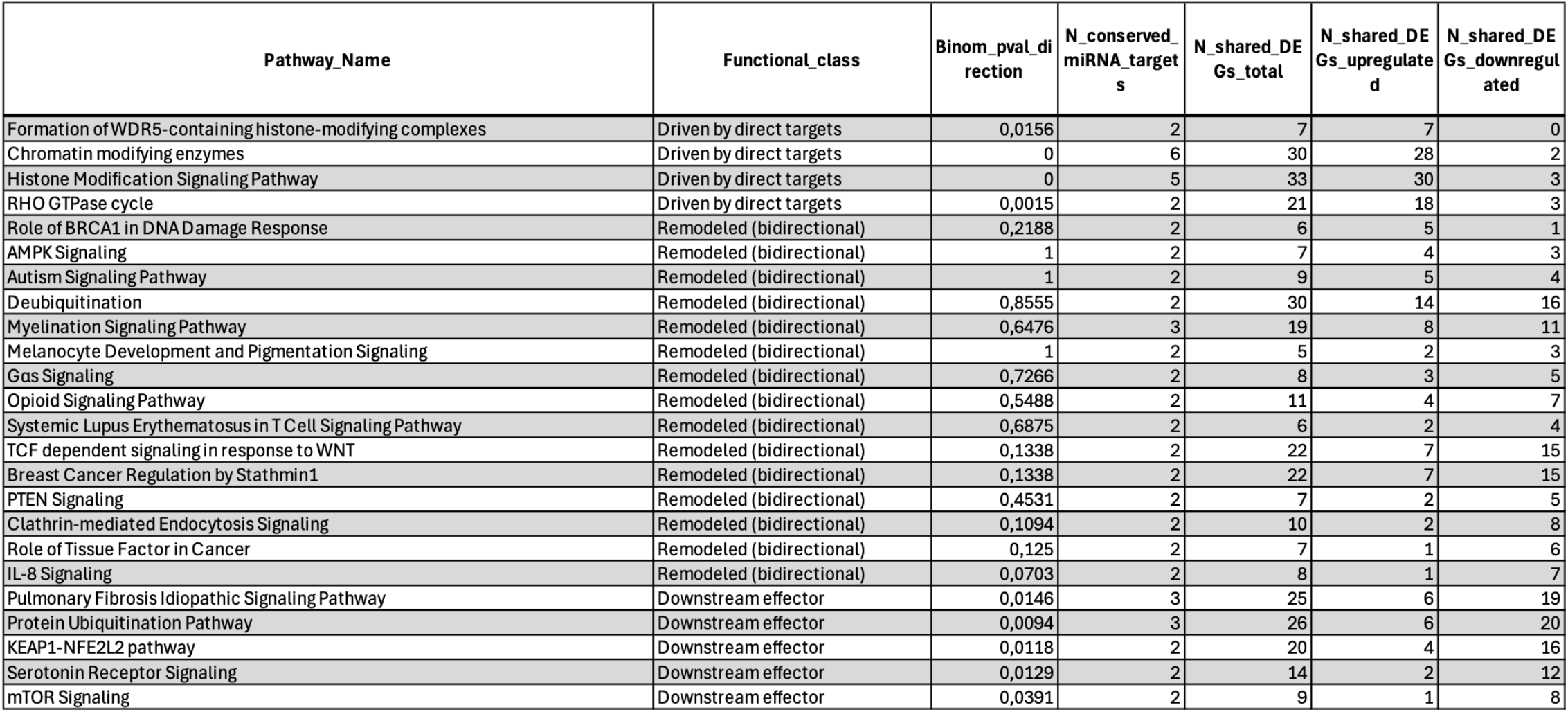

